# Three PilZ domain proteins, PlpA, PixA and PixB, have distinct functions in regulation of motility and development in *Myxococcus xanthus*

**DOI:** 10.1101/2021.03.04.433885

**Authors:** Sofya Kuzmich, Dorota Skotnicka, Dobromir Szadkowski, Philipp Klos, María Pérez-Burgos, Eugenia Schander, Dominik Schumacher, Lotte Søgaard-Andersen

## Abstract

In bacteria, the nucleotide-based second messenger bis-(3’-5’)-cyclic dimeric GMP (c-di-GMP) binds to effectors to generate outputs in response to changes in the environment. In *Myxococcus xanthus*, c-di-GMP regulates type IV pili-dependent motility and the starvation-induced developmental program that results in the formation of spore-filled fruiting bodies; however, little is known about the effectors that bind c-di-GMP. Here, we systematically inactivated all 24 genes encoding PilZ domain-containing proteins, which are among the most common c-di-GMP receptors. We confirm that PlpA, a stand-alone PilZ-domain protein, is specifically important for motility and that Pkn1, which is composed of a Ser/Thr domain and a PilZ domain, is specifically important for development. Moreover, we identify two PilZ-domain proteins that have distinct functions in regulating motility and development. PixB, which is composed of two PilZ domains and an acetyltransferase domain, binds c-di-GMP *in vitro* and regulates type IV pili-dependent and gliding motility upstream of the Frz chemosensory system as well as development. The acetyltransferase domain is required and sufficient for function during growth while all three domains and c-di-GMP binding are essential for PixB function during development. PixA is a response regulator composed of a PilZ domain and a receiver domain, binds c-di-GMP *in vitro*, and regulates motility downstream of the Frz chemosensory system by setting up the polarity of the two motility systems. Our results support a model whereby the three proteins PlpA, PixA and PixB act in parallel pathways and have distinct functions to regulation of motility.

**Importance:** c-di-GMP signaling controls bacterial motility in many bacterial species by binding to downstream effector proteins. Here, we identify two PilZ domain-containing proteins in *Myxococcus xanthus* that bind c-di-GMP. We show that PixB, which contains two PilZ domains and an acetyltransferase domain, acts upstream of the Frz chemosensory system to regulate motility via the acetyltransferase domain while the intact protein and c-di-GMP binding are essential for PixB to support development. By contrast, PixA acts downstream of the Frz system to regulate motility. Together with previous observations, we conclude that PilZ-domain proteins and c-di-GMP act in multiple parallel pathways to regulate motility and development in *M. xanthus*.

## Introduction

Bacteria have evolved different strategies that allow them to sense and subsequently adapt and differentiate in response to changing environmental conditions. One strategy centers on signaling by a variety of nucleotide-based second messengers (1). The highly versatile second messenger bis-(3’-5’)-cyclic dimeric GMP (c-di-GMP) is found widespread in bacteria and is involved in many aspects of bacterial physiology including regulation of motility, adhesion, synthesis of secreted polysaccharides, biofilm formation, cell cycle progression, development, and virulence (2, 3).

Signaling by c-di-GMP depends on its regulated synthesis by diguanylate cyclases (DGCs), which contain an enzymatically active GGDEF domain, and degradation by phosphodiesterases (PDEs), which contain either a catalytic EAL or HD-GYP domain (2, 3). Output responses are generated by binding of c-di-GMP to and allosteric regulation of effectors, which direct downstream responses at the transcriptional, translational or post-translational level (2, 3). Several different protein effectors with little sequence homology have been identified and includes enzymatically inactive GGDEF and EAL domain proteins (4–8), PilZ-domain proteins (9–15), MshEN-domain proteins (16, 17), members of different transcription factors families as well as a nucleoid-associated DNA binding protein (18–26), and ATPases of flagella, type III and type VI secretion systems (27). The number of DGCs and PDEs encoded by many bacterial genomes typically outnumbers that of known c-di-GMP effectors (2, 28), thus hampering a detailed understanding of how effects of changing c-di-GMP levels are implemented.

*Myxococcus xanthus*, a Gram-negative deltaproteobacterium, is a model organism for studying how social behaviors in bacteria can be modulated by environmental cues (29, 30). In the presence of nutrients, *M. xanthus* cells grow, divide and actively move by means of two motility systems to generate colonies in which cells spread outward in a highly coordinated fashion and prey on other microorganisms if present. In response to nutrient depletion, a developmental program is initiated that results in the formation of multicellular, spore-filled fruiting bodies. Both social behaviors depend on regulated motility (31, 32) as well as signaling by c-di-GMP (33).

The rod-shaped *M. xanthus* cells move on surfaces in the direction of their long axis using two polarized motility systems and with clearly defined leading and lagging cell poles (31, 32). Gliding motility is favorable on hard and dry surfaces (34) and generally allows the movement of single cells (35). Gliding depends on the Agl/Glt complexes, which assemble at the leading cell pole, attach to the substratum, and disassemble as they reach the lagging cell pole (36, 37). By contrast, type IV pili (T4P)-dependent motility is favorable on soft and moist surfaces and mostly occurs in groups of cells (34, 35). T4P localize to the leading cell pole and undergo cycles of extension, surface adhesion, and retraction to pull cells across surfaces (38–40). Moreover, T4P-dependent motility in *M. xanthus* depends on a secreted exopolysaccharide (EPS) (41–45). The characteristic polarized assembly of the two motility machineries at the leading cell pole only is regulated by a set of four proteins that make up the so-called cell polarity module. The key protein in this module is the small GTPase MglA, which in its GTP-bound form localizes at the leading cell pole to stimulate assembly of the Agl/Glt complexes as well as formation of T4P (36, 46–49). The activity and localization of MglA is regulated by its cognate Guanine nucleotide Exchange Factor (GEF) composed of the RomR and RomX proteins (50) and its cognate GTPase Activating Protein (GAP) MglB (46, 49). These three proteins localize in bipolar, asymmetric patterns to the cell poles and with the GAP activity dominating at the lagging cell pole and the GEF activity dominating at the leading cell pole (46, 49–51). Importantly, the MglB GAP activity at the lagging pole blocks MglA-GTP accumulation at this pole and therefore ensures that T4P only assemble at the leading pole (48). Similarly, because MglA-GTP is incorporated into the Agl/Glt complexes, these complexes disassemble as their reach the lagging pole (36).

Consequently, cells lacking MglB undergo frequent so-called pseudo-reversals because they have T4P at both poles and do not disassemble the Agl/Glt complexes at the lagging pole (36, 48). *M. xanthus* cells also undergo reversals in a regulated manner stimulated by the Frz chemosensory system (52). The two output response regulators of this system, FrzX (53) and FrzZ (54, 55), induce the relocalization of MglA, MglB, RomR and RomX between the two poles (46, 49–51), thus laying the foundation for T4P formation and Agl/Glt complex assembly at the new leading cell pole after a reversal. c-di-GMP accumulates during growth of *M. xanthus* and at a 10-fold higher level during development (56, 57). During growth, c-di-GMP regulates T4P-dependent motility by regulating transcription of the *pilA* gene, which encodes the major pilin subunit of T4P, and EPS synthesis (7, 56). During development, the increased c-di-GMP level induces an increase in EPS synthesis that is essential for fruiting body formation and sporulation (57). Among the 17 GGDEF domain containing proteins encoded by the *M. xanthus* genome, only DmxA and DmxB have been experimentally shown to have DGC activity (56, 57). Lack of DmxA only causes defects during growth and is important for T4P-dependent motility (56).

The c-di-GMP receptors involved in regulating *pilA* transcription and EPS synthesis during growth are not known. Lack of DmxB only causes defects during development, and DmxB is responsible for the increase in the c-di-GMP level during development (57). c-di-GMP binds to the transcriptional regulator EpsI/Nla24 (57), which is an enhancer binding protein important for expression of genes encoding proteins for EPS synthesis (58). It was suggested that c-di-GMP may bind to EpsI/Nla24 during development to stimulate EPS synthesis (57). However, generally, it is largely unknown how effects of changing c-di-GMP levels are implemented to affect motility and development.

Here, we aimed to further understand the molecular basis of how the effects of changing levels of c-di-GMP are implemented in *M. xanthus*. Because PilZ-domain proteins in several species have been shown to be involved in regulation of flagella-based motility (14, 59–63), T4P-dependent motility (64, 65), and synthesis of secreted polysaccharides (12, 66, 67), we addressed the function of PilZ-domain proteins in *M. xanthus*. Previously, bioinformatics analyses identified 24 PilZ-domain proteins in *M. xanthus* (9), among which only three have been analyzed in some details experimentally (see below). Here, we report the identification of two PilZ-domain proteins, renamed to PixA (MXAN_1087/MXAN_RS05220 (old/new annotation)) and PixB (MXAN_2604/MXAN_RS12590), that bind c-di-GMP *in vitro*. PixB is composed of two PilZ domains and an acetyltransferase domain. PixB functions upstream of the Frz system to regulate reversals and is also important for development. The acetyltransferase activity is essential for activity and, likely stimulated by c-di-GMP binding to the PilZ domains during development. PixA is also important for a correct reversal frequency; however, PixA acts downstream of the Frz system to regulate the polarity of the motility systems and lack of PixA causes pseudo-reversals.

## Results

### Phenotypic characterization of mutants lacking individual PilZ domain proteins

The ∼110 amino acid PilZ domain is widely distributed in bacteria and can be found either as a stand-alone domain in PilZ single domain proteins or in combination with other domains (9). Genome analyses previously revealed that the *M. xanthus* genome encodes 24 proteins with a PilZ-domain (9) (Fig. 1; S1). 14 are composed only of a PilZ domain while the remaining contain additional domains. Some of the latter are typically involved in signal transduction while four proteins contain a DnaK or DnaJ domain suggesting that they might be involved in protein folding and/or quality control. In fully sequenced Myxococcales genomes, the 24 PilZ-domain proteins are largely conserved in closely related fruiting Cystobacterineae but not in the non-fruiting Cystobacterineae and the more distantly related fruiting Nannocystineae and Sorangineae (Fig. S2A).

**Figure 1.**
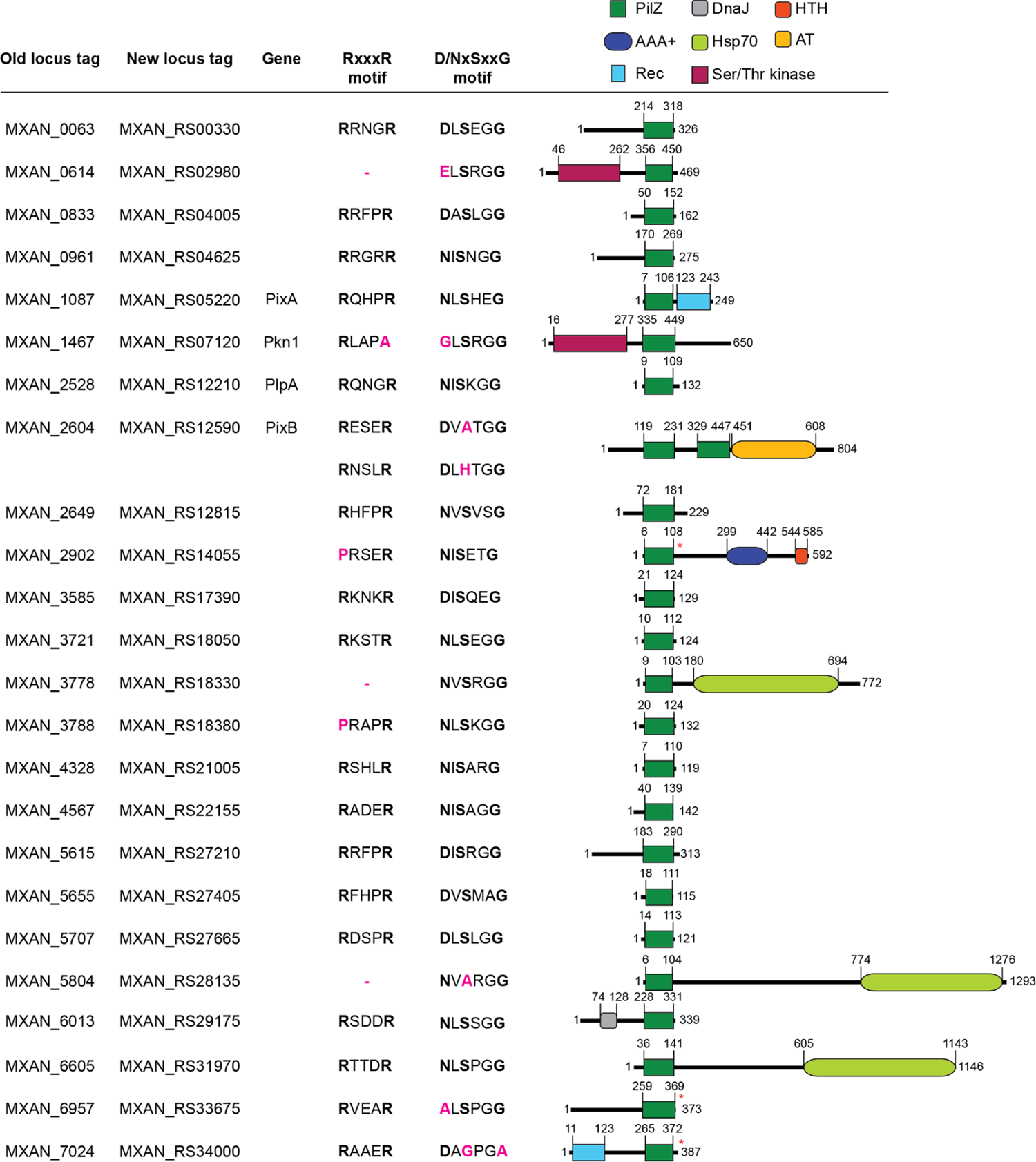
Analysis of *M. xanthus* proteins containing a PilZ domain. Domain organization of *M. xanthus* proteins containing a PilZ domain. Locus tags according to the original MXAN annotation of the *M. xanthus* genome and the most recent annotation are included together with names of proteins described in the literature (See text). Protein domains were identified with Pfam, except the PilZ domains marked with a red asterisk, which were identified with Motif Scan (See Materials & Methods). Domains are drawn to scale. The two conserved sequence motifs involved in c-di-GMP binding are indicated at the top (2); the corresponding amino acid residues are indicated in black if conserved and pink for non-conserved residues.

Among the 24 PilZ domain proteins, three have been previously characterized experimentally. Pkn1 is a Ser/Thr kinase that is important for development (68) (Fig. 1). Based on sequence analysis, the PilZ domain of Pkn1 is predicted not to bind c-di-GMP; however, c-di-GMP binding by Pkn1 has not been studied. PlpA is a single domain PilZ domain protein (Fig. 1), which is important for regulation of the reversal frequency, but not for development (69). Despite possessing the conserved residues in its PilZ domain for c-di-GMP binding (Fig. 1), PlpA was reported not to bind c-di-GMP *in vitro* and substitutions of amino acids predicted to be important for c-di-GMP binding did not cause reversal defects *in vivo* (69). MXAN_2902 is an enhancer binding protein (Fig. 1), which is important for fruiting body morphology under a subset of starvation conditions (70) and contains a PilZ domain that is predicted not to bind c-di-GMP; however, c-di-GMP binding has not been studied.

We systematically generated in-frame deletion mutations in all 24 genes encoding PilZ domain proteins and tested the strains for motility, EPS accumulation (Fig. 2) and development (Fig. 3). On 0.5% agar, which is favorable for T4P-dependent motility, the wild-type (WT) strain DK1622 formed the long flares characteristic of T4P-dependent motility while the Δ*pilA* mutant, which lacks the major pilin of T4P, did not (Fig. 2). Among the 24 mutants generated, the Δ*plpA*, ΔMXAN_1087 and ΔMXAN_2604 had a defect in T4P-dependent motility with the formation of shorter flares and significantly reduced colony expansion. Because EPS is important for T4P-dependent motility and its synthesis is regulated by c-di-GMP, we determined EPS synthesis using a Trypan Blue-based colorimetric assay, and observed that all 24 mutants synthesized EPS as WT while the level was decreased in the Δ*pilA* mutant, which served as a negative control (71) (Fig. 2). On 1.5% agar, which is favorable to gliding motility, WT displayed the single cell movement at the edge of the colony, whereas the Δ*aglQ* mutant, which lacks the motor of the gliding motility complex (72, 73), did not (Fig. 2). Among the 24 mutants generated, the Δ*plpA*, ΔMXAN_1087 and ΔMXAN_2604 mutants had a defect in gliding with fewer single cells at the colony edge and significantly reduced colony expansion. These observations are in agreement with previous analyses of a *plpA* mutant (69).

**Figure 2.**
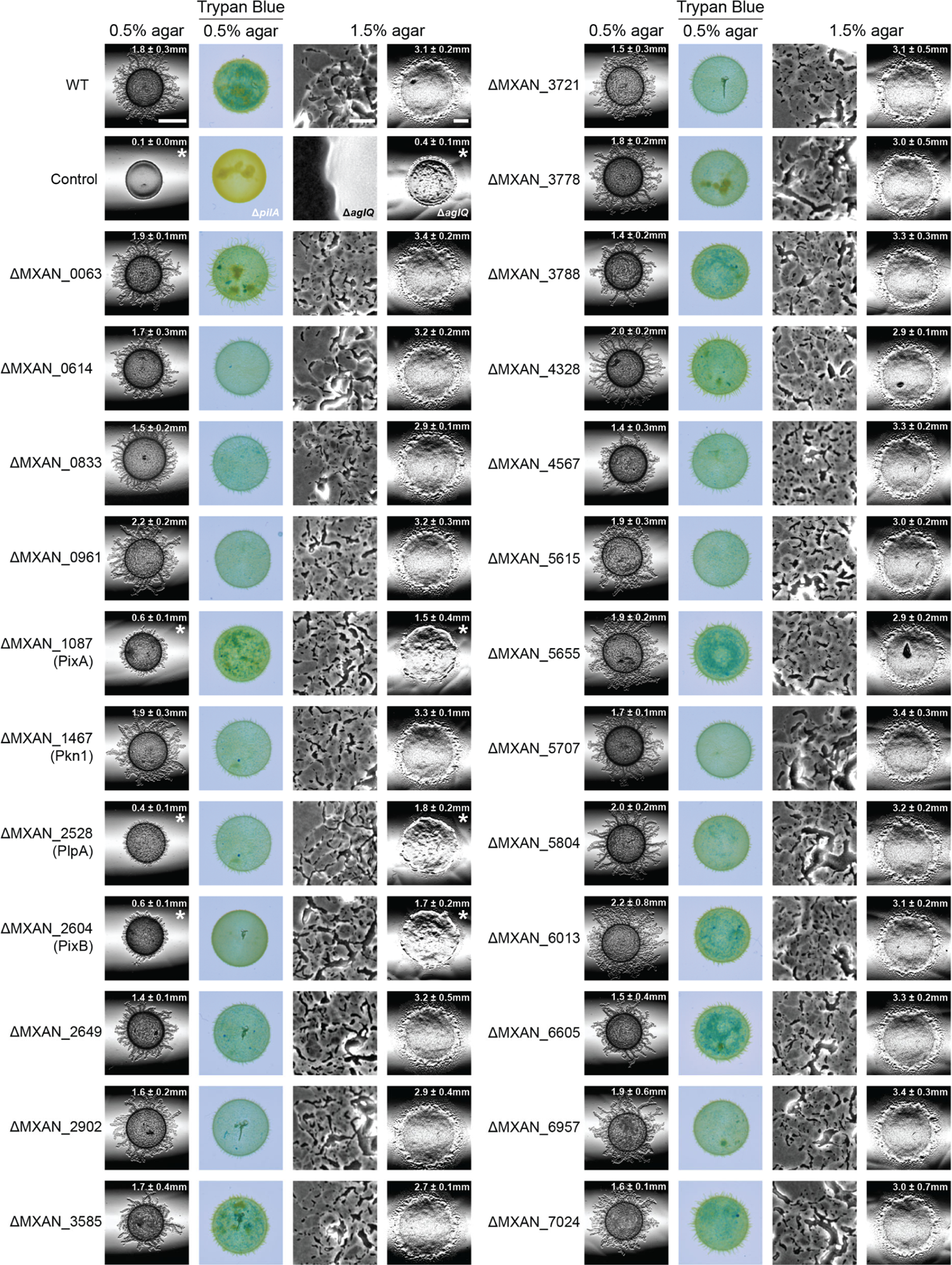
Motility assays for mutants lacking individual PilZ domain proteins. T4P-dependent motility and gliding motility were analyzed on 0.5% agar and 1.5% agar supplemented with 0.5% CTT, respectively. Motility was quantified by the increase in colony radius and numbers indicate the mean increase in colony radius ± standard deviation (SD) from three biological replicates after 24 h. *, *P* < 0.05 in a Student’s t test. Scale bars, 3 mm (0.5% agar), 100 μm (1.5% agar, left), 3 mm (1.5% agar, right). EPS accumulation was determined on 0.5% agar supplemented with 0.5% CTT and 10 μg/ml Trypan Blue.

**Figure 3.**
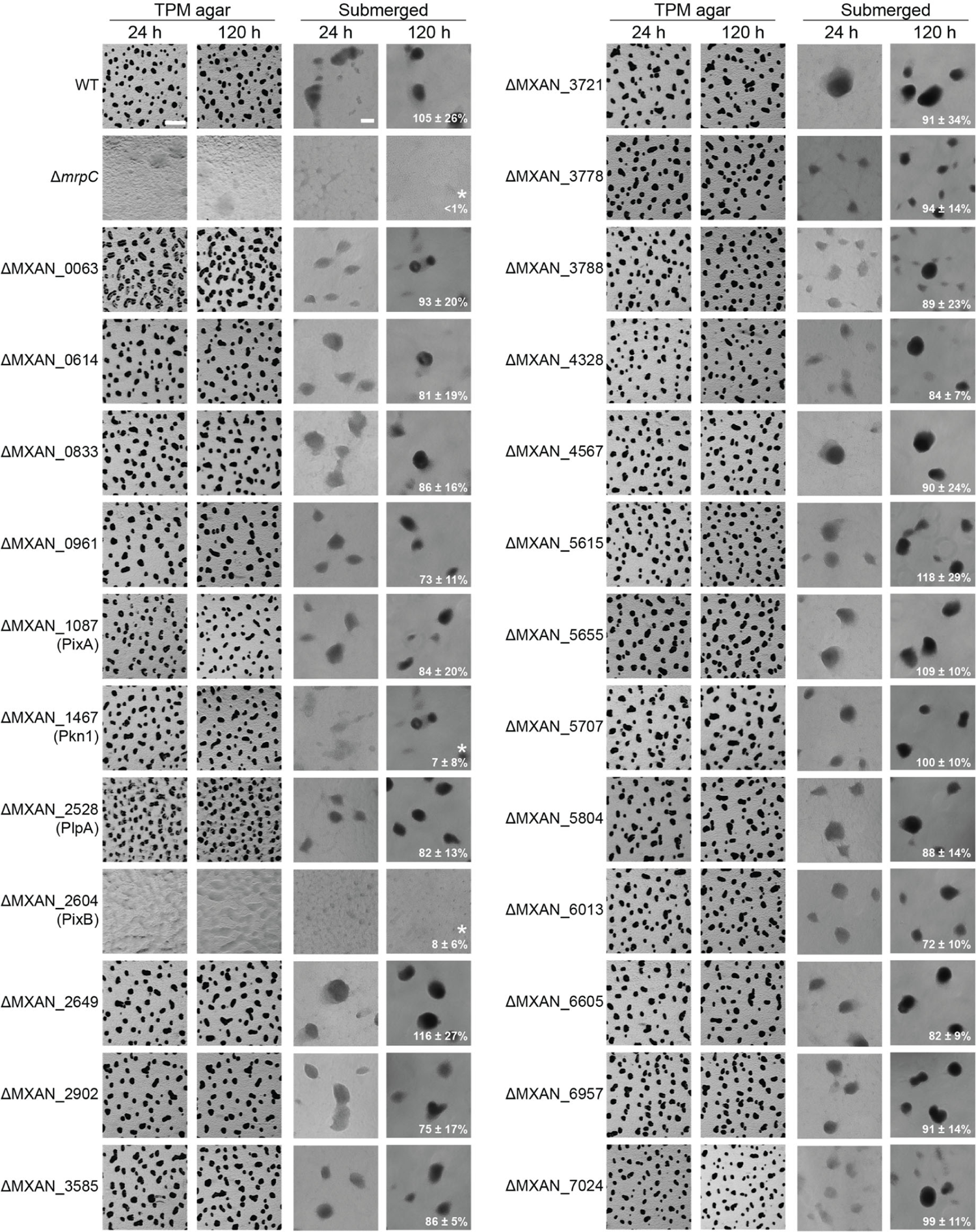
Development assays for mutants lacking individual PilZ domain proteins. Fruiting body formation and sporulation on TPM agar and in MC7 submerged culture. Numbers indicate heat- and sonication resistant spores formed at 120 h of starvation in submerged culture in percentage of WT (100%) ± SD from three biological replicates. *, *P* < 0.05 in Student’s t test. Scale bars, 500 µm (TPM agar), 100 µm (submerged).

We tested the ability of the 24 generated mutants to undergo development with fruiting body formation and sporulation by starving cells under to different condition, i.e. on TPM agar and on a polystyrene surface under submerged conditions. Under both conditions, WT cells had formed fruiting bodies at 24 h and spores at 120 h as measured for cells starved under submerged conditions, while the Δ*mrpC* mutant, which lacks a transcription factor important for development and served as a negative control (74), had not (Fig. 3). Among the 24 mutants generated, only the Δ*pkn1* and ΔMXAN_2604 mutants had developmental defects. In agreement with previous observations (68), the Δ*pkn1* mutant formed fruiting bodies but was reduced in sporulation; the ΔMXAN_2604 mutant did not form fruiting bodies under any of the two conditions tested and was also reduced in sporulation. We did not observe developmental defects in the ΔMXAN_2902 mutant under the two conditions tested (Fig. 3).

We conclude that the Δ*plpA*, ΔMXAN_1087 and ΔMXAN_2604 mutations cause defects in both motility systems and the Δ*pkn1* and ΔMXAN_2604 mutations cause defects in development. From here on, we focused on MXAN_1087 and MXAN_2604 that we renamed PixA (MXAN_1087/MXAN_RS05220 (old/new annotation)) and PixB (MXAN_2604/MXAN_RS12590/) for <u>P</u>ilZ-domain protein in *M. <u>x</u>anthus* <u>A</u> and <u>B</u>.

### PixA and PixB have distinct functions in regulation of motility

PixA is a response regulator of two-component systems with an N-terminal PilZ domain that possesses all necessary residues for c-di-GMP binding (Fig. 1; Fig. S1) and a C-terminal receiver domain in which the conserved residues important for phosphorylation including the potentially phosphorylatable Asp180 residue are conserved (Fig. S3) (75). PixB consists of two PilZ domains that both contain the conserved RxxxR motif for c-di-GMP binding but lack the conserved Ser residue in the second motif (Fig. 1; Fig. S1), and an acetyltransferase (AT) domain of the Gcn5-related *N*-acetyltransferase (GNAT) family. Both proteins as well as the genetic neighborhood of the respective genes are conserved in closely related fruiting Cystobacterineae (Fig. S2A-C).

The distance between *pixA* and the downstream gene supports that *pixA* is not part of an operon while a similar analysis of the *pixB* locus supports that *pixB* could be in an operon with the two downstream genes (Fig. S2BC). To determine whether *pixA* and *pixB* are indeed important for motility and development in the case of *pixB*, we ectopically expressed the full-length genes under the control of their native promoter (P_nat_)from the *attB* site in the Δ*pixA* and Δ*pixB* mutants (Fig. S2BC). Motility assays on 0.5% and 1.5% agar plates revealed that the motility defects of both mutants were fully corrected in the two complementation strains (Fig. 4A); similarly, the developmental defects of the Δ*pixB* mutant were complemented by ectopic expression of *pixB* (Fig. 4B). Because the increased c-di-GMP level during development is important for EPS synthesis (57), we determined EPS accumulation in developing cells. We observed that the Δ*pixB* mutant accumulated EPS similarly to WT as estimated by the Trypan Blue binding assay (Fig. 4B).

**Figure 4.**
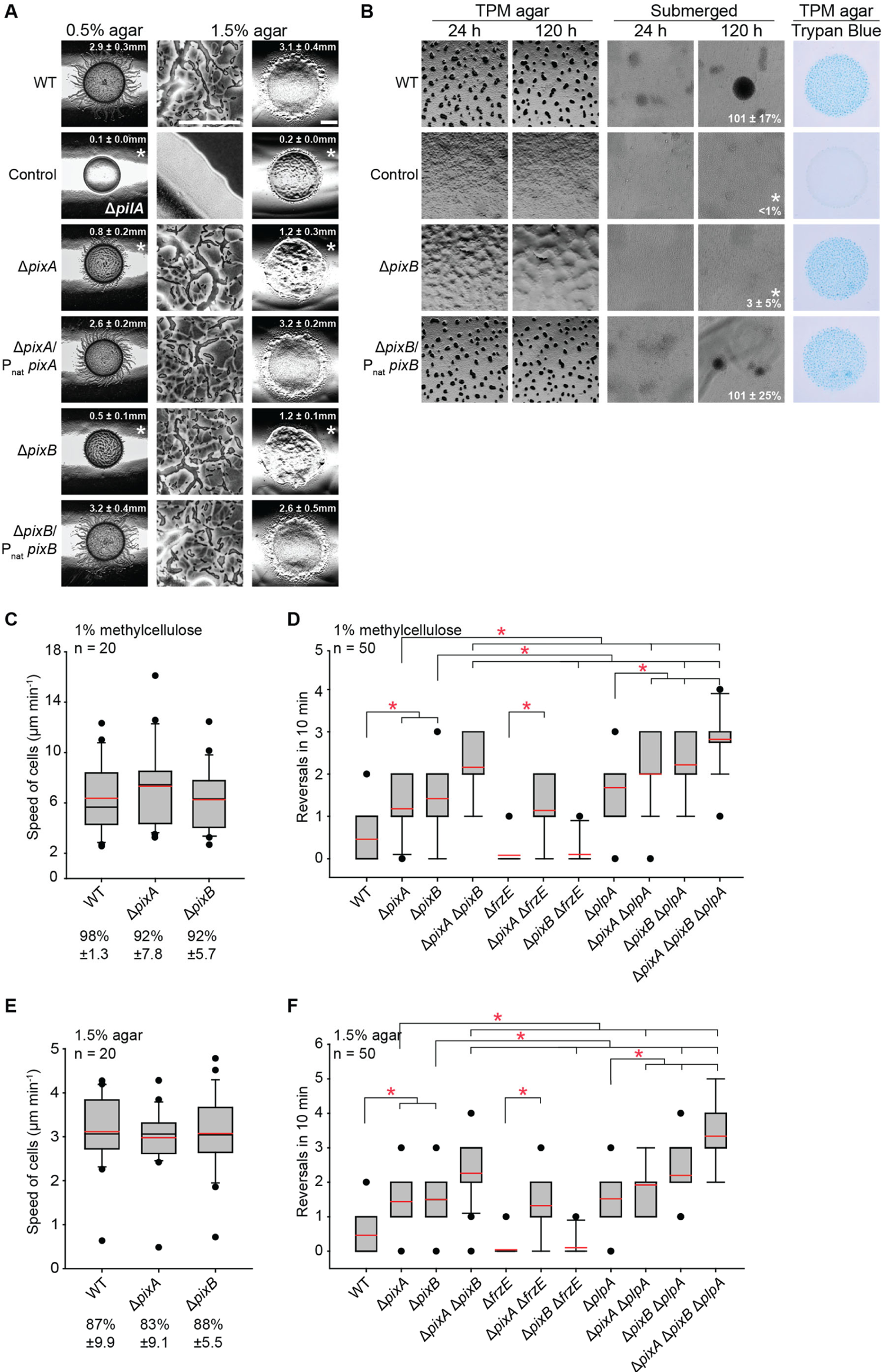
PixA is important for motility and PixB for motility and development A. Complementation of Δ*pixA and* Δ*pixB* motility defects. Assays were done as in Fig. 2. Scale bars, 2 mm (0.5% agar), 100 μm (1.5% agar, left), 3 mm (1.5% agar, right). B. Complementation of Δ*pixB* developmental defects. Fruiting body formation and sporulation were analyzed as in Fig. 3. EPS accumulation was determined on TPM agar with addition of Trypan Blue. Numbers indicate heat- and sonication resistant spores formed at 120 h of starvation in submerged culture in percentage of WT (100%) ± SD from three biological replicates. *, *P* < 0.05 in Student’s t test. Scale bars, 500 µm (TPM agar), 100 µm (submerged). C, E. Speed of Δ*pixA* and Δ*pixB* cells in 1% methylcellulose (C) and on 1.5% agar supplemented with 0.5% CTT (E). Boxplots represent the speed of isolated cells per min, boxes enclose the 25^th^ and 75^th^ percentile, whiskers represent the 10^th^ and 90^th^ percentile and dots outliers, red and black line indicates the mean and median, respectively. * *P* < 0.05 in Mann-Whitney Rank Sum Test, n = 20 cells. Numbers below indicate the fraction of cells that displayed single cell movement ± SD. D, F. Reversal frequency of Δ*pixA*, Δ*pixB* and Δ*plpA* mutants on 1% methylcellulose (D) and 1.5% agar (F). Boxplots represent the reversals of cells per 10 min. Boxplots are as in C and E. * *P* < 0.05 in Mann-Whitney Rank Sum Test, n = 50 cells.

Motility defects scored in a population-based assay can be caused by *bona fide* motility defects or an altered reversal frequency. To discriminate between these defects, we scored the speed and reversal frequency of single cells moving preferentially by T4P-dependent motility in 1% methylcellulose and by gliding on 1.5% agar in single cell motility assays. Δ*pixA* and Δ*pixB* cells moved with the same speed as WT cells under both conditions (Fig. 4CE) but reversed significantly more frequently than WT under both conditions (Fig. 4DF).

To test whether PixA and PixB regulate cellular reversals in a Frz-dependent manner, we deleted *frzE*, which encodes the FrzE kinase of the Frz system (76), in the Δ*pixA* and Δ*pixB* mutants. The Δ*frzE* mutant had a hypo-reversing phenotype under both conditions tested while the Δ*frzE* Δ*pixA* hyper-reversed and the Δ*frzE* Δ*pixB* mutant, similarly to the Δ*frzE* mutant, hypo-reversed (Fig. 4DF). These observations suggest that PixA acts downstream of the Frz system to inhibit reversals while PixB acts upstream of the Frz system to inhibit reversals.

Lack of PlpA also causes cells to hyper-reverse and PlpA has been suggested to act downstream of the Frz system to inhibit reversals (69). We used epistasis experiments to address whether PixA, PixB and PlpA act in the same pathway. We confirmed that the Δ*plpA* mutant hyper-reverses; more importantly, all three double mutants had an additive phenotype and reversed more frequently than the three single mutants (Fig. 4DF). Finally, the triple mutant reversed even more frequently than the three double mutants (Fig. 4DF). Altogether, these observations demonstrate that PixA and PixB, similarly to PlpA, are not important for motility *per se* but for cells to move with the correct reversal frequency.

Moreover, the epistasis tests support that these three proteins act in independent pathways to establish the correct reversal frequency, and with PixB acting upstream of the Frz system to inhibit reversals while PixA and PlpA act downstream of the Frz system to inhibit reversals.

### PixA is important for unipolar localization of MglA

Reversals are induced by Frz signaling while pseudo-reversals, which occur independently of the Frz system, are caused by interfering with the correct localization of the four proteins of the polarity module. In particular, mutations that cause bipolar localization of MglA also cause hyper-reversals (46, 49, 77). Because lack of PixA causes hyper-reversals independently of the Frz system (Fig. 4DF), i.e. lack of PixA causes pseudo-reversals, we hypothesized that PixA would be important for correct MglA localization. To this end, we expressed an MglA-mVenus fusion protein from the native site in WT and the Δ*pixA* mutant. Epi-fluorescence microscopy demonstrated that MglA-mVenus in the absence of PixA localizes in a more symmetric bipolar pattern than in WT while the total polar signal remained unchanged (Fig. 5). Lack of PlpA also causes a shift in MglA-GTP localization towards symmetric bipolar (69). Altogether, these observations together with the epistasis analyzes demonstrate that PixA and PlpA act in independent pathways to set up the correct localization pattern of the proteins of the polarity module and, in particular, to establish a unipolar localization of MglA.

**Figure 5.**
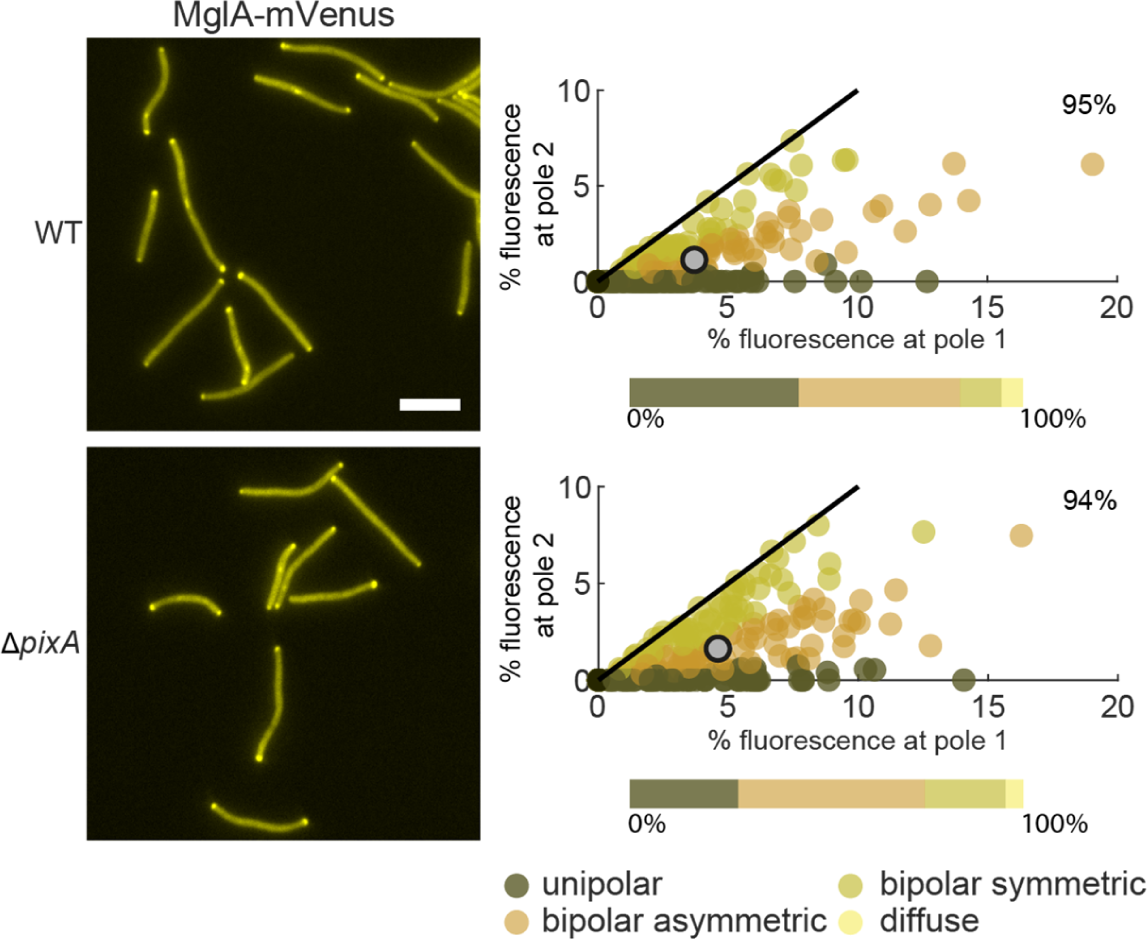
PixA is important for unipolar MglA-mVenus localization For each cell with polar clusters, an asymmetry index was calculated as described in Materials & Methods to distinguish between unipolar, asymmetric bipolar and symmetric bipolar localization. Cells with no polar signal were categorized as cells with diffuse localization. In the scatter plot, the percentage of total fluorescence at pole 2 is plotted against the percentage of total fluorescence at pole 1 for all cells with polar cluster(s) and individual cells are color-coded according to its localization pattern. Pole 1 is per definition the pole with the highest fluorescence. Black lines are symmetry lines, grey dots show the mean and numbers in the upper right corner are the mean percentage of total fluorescence in the cytoplasm. Horizontal bars show the percentage of cells with polar localization patterns and diffuse localization according to the color code. n = 200 cells. Scale bar, 5 µm.

### PixA and PixB bind c-di-GMP *in vitro*

The PilZ domain in PixA contains the two motifs important for c-di-GMP binding while both PilZ domains in PixB have a substitution in the most C-terminal of the two motifs (Fig. 1; Fig. S1). To determine whether PixA and PixB are can bind c-di-GMP, we performed a differential radial capillary action of ligand assay (DRaCALA) using purified full-length PixA-His_6_ and PixB-His_6_ (Fig. 6AB; Fig. S4). PixA-His_6_ and PixB-His_6_ specifically bound ^32^P-labeled c-di-GMP (Fig. 6AB). As expected, the PixA^R9A^-His_6_ variant, which contains the Arg9 to Ala substitution in the N-terminal part of the bipartite c-di-GMP binding motif (Fig. 6A; Fig. S4), did not detectably bind ^32^P-c-di-GMP (Fig. 6A). To determine which of the PilZ domains in PixB are involved in c-di-GMP binding, we overexpressed the PixB^R121A^-His_6_, PixB^R331A^-His_6_ and PixB^R121A/R331A^-His_6_ variants, which contain a substitution in the N-terminal part of the c-di-GMP binding motif (Fig. 6B). However, we were not able to purify any of these variants because they formed inclusion bodies under all conditions tested.

**Figure 6.**
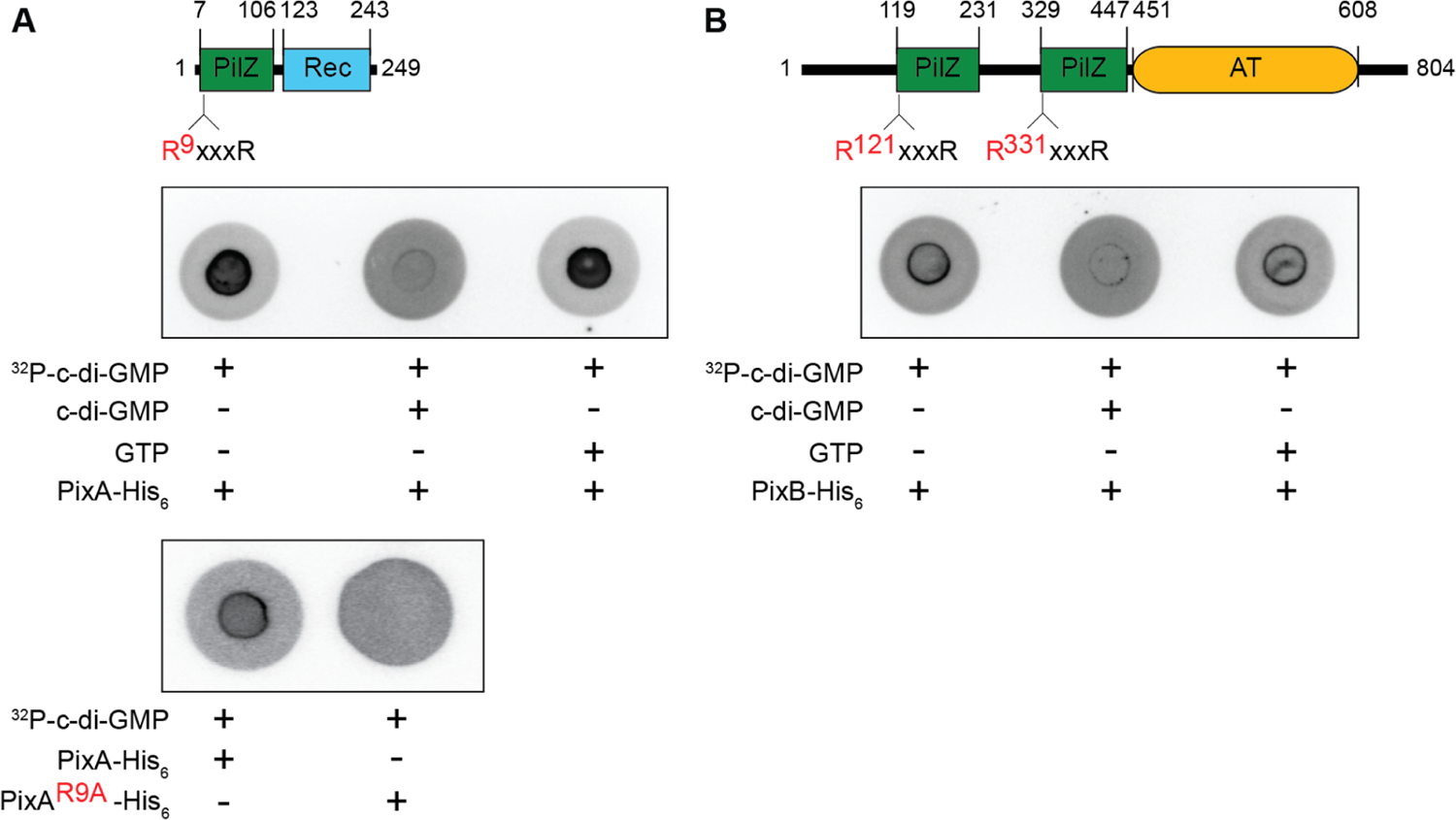
*In vitro* assay for c-di-GMP binding by PixA and PixB DRaCALA assay to detect specific c-di-GMP binding by purified proteins. Full-length PixA-His_6_ (A, upper panel), PixA^R9A^-His_6_ (A, lower panel) or PixB-His_6_ (B) were incubated at a final concentration of 20 µM with ^32^P-labeled c-di-GMP. Unlabeled c-di-GMP and GTP were added to final concentrations of 400 µM, as indicated.

We conclude that PixA binds c-di-GMP *in vitro* and that the R^9^xxxR motif is important for this binding; PixB also binds c-di-GMP *in vitro* but it is unclear which domain(s) is involved.

### Role of c-di-GMP binding by PixA and PixB *in vivo*

To test whether c-di-GMP binding is important for PixA and PixB function *in vivo*, we ectopically expressed FLAG-tagged full-length WT and mutant variants from the Mx8 *attB* site in the relevant in-frame deletion mutants (Fig. 7AB). PixA^WT^-FLAG and PixA^R9A^-FLAG accumulated at similar levels during growth (Fig. 7C) and restored motility including the reversal frequency in the Δ*pixA* mutant (Fig. 7A; Fig. S5AB).

**Figure 7.**
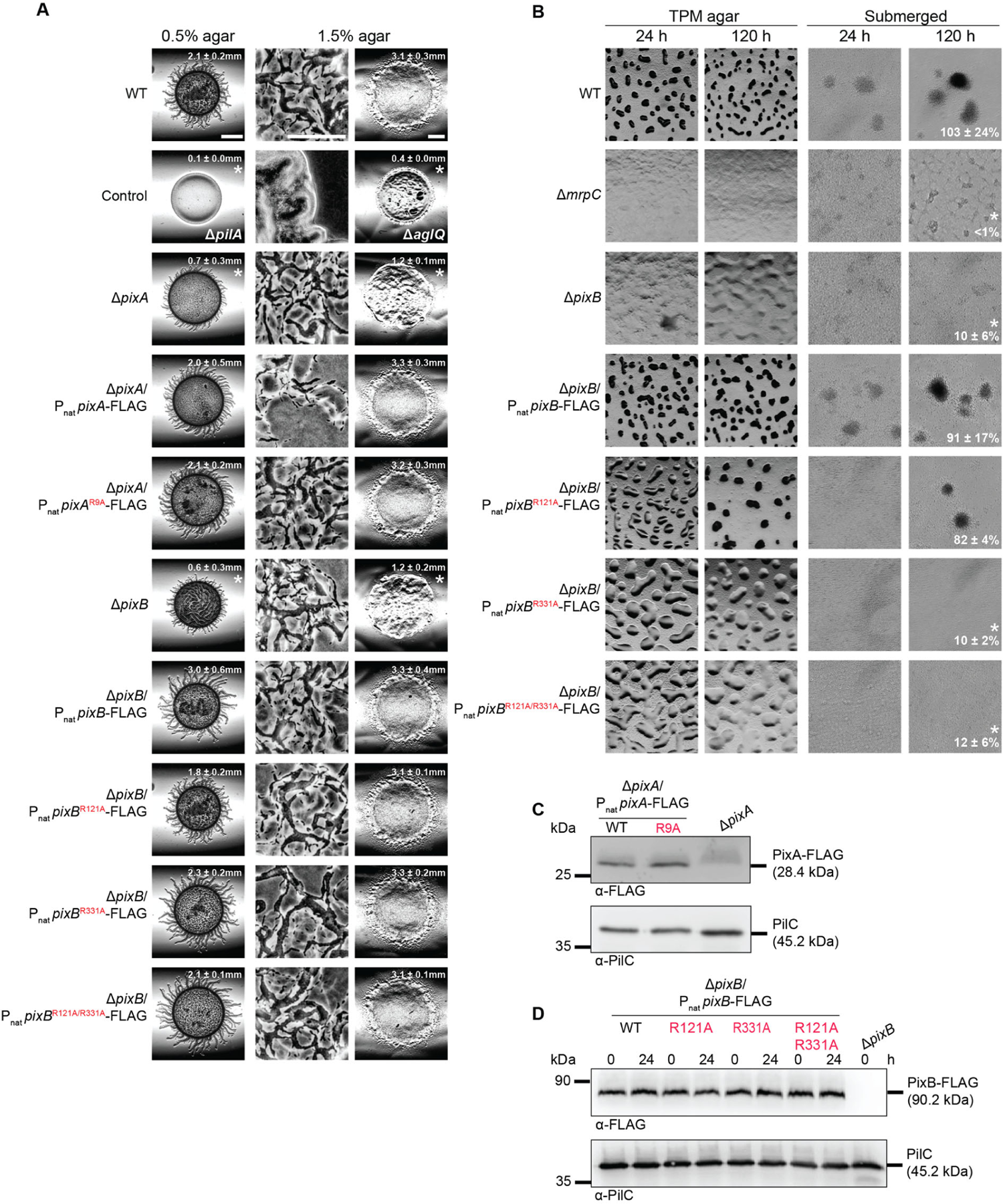
Role of c-di-GMP binding by the PixA and PixB PilZ domains in regulation of motility and development A. Motility assays of strains expressing full-length WT or protein variants with a mutated RxxxR motif in the PilZ domains. Motility was analyzed as described in Fig. 2. Scale bars, 3 mm (0.5% agar), 100 μm (1.5% agar, left), 3 mm (1.5% agar, right). B. Development assays of the Δ*pixB* mutant expressing full-length WT or protein variants with a mutated RxxxR motif in PilZ domains. Assays were done as described in Fig. 3. Scale bars, 500 µm (TPM agar), 100 µm (submerged). C, D. Immunoblot detection of PixA-FLAG (C) and PixB-FLAG (D). For PixA-FLAG samples, total cell extracts were prepared form exponentially grown cells; for PixB-FLAG samples, cell extracts were prepared form exponentially grown cells and at 24 h of starvation in submerged culture. 10 µg of protein was loaded per lane and samples separated by SDS-PAGE. Upper blots were probed with α-FLAG antibodies and lower blots with α-PilC antibodies. The PilC blots served as loading controls. Molecular mass marker is indicated on the left.

PixB^WT^-FLAG and the variants PixB^R121A^-FLAG, PixB^R331A^-FLAG and PixB^R121A/R331A^-FLAG accumulated at similar levels during growth and development and all four proteins restored the motility defects of the Δ*pixB* mutant (Fig 7AD; Fig. S5AB). While fruiting body formation and sporulation was fully restored by PixB^WT^-FLAG (Fig. 7B), the strain complemented with PixB^R121A^-FLAG displayed delayed fruiting body formation on both solid and submerged condition but restored sporulation at 120 h; however, PixB^R331A^-FLAG and PixB^R121A/R331A^-FLAG neither restored fruiting body formation nor sporulation in the Δ*pixB* mutant (Fig. 7B).

These observations support that c-di-GMP binding by PixA and PixB under the conditions tested is not important for their function in motility. By contrast, c-di-GMP binding by the two PilZ domains in PixB appears to be important for fruiting body formation and sporulation, and with the C-terminal domain being the most important.

### Structure-function analysis of PixA and PixB

The receiver domain of PixA contains the conserved residues important for phosphorylation including the putative phosphorylated residue Asp180 (Fig. S3). Substitution of the phosphorylatable Asp to Glu mimics the phosphorylated form in some response regulators (78, 79), while the Asp to Asn substitution blocks phosphorylation. Therefore, to test whether phosphorylation of the receiver domain could be important for PixA function, we ectopically expressed FLAG-tagged full-length PixA variants with a D180E or D180N substitution. Only PixA^WT^-FLAG and PixA^D180N^-FLAG restored motility including the reversal frequency in the Δ*pixA* mutant while PixA^D180E^-FLAG did not (Fig. 8A; Fig. S5CD). PixA^WT^-FLAG, PixA^D180E^-FLAG and PixA^D180N^-FLAG accumulated at similar levels during growth (Fig. 8B). Altogether, these findings support a scenario whereby PixA might be phosphorylated and in which the non-phosphorylated form of PixA would represents the active form of the protein.

**Figure 8.**
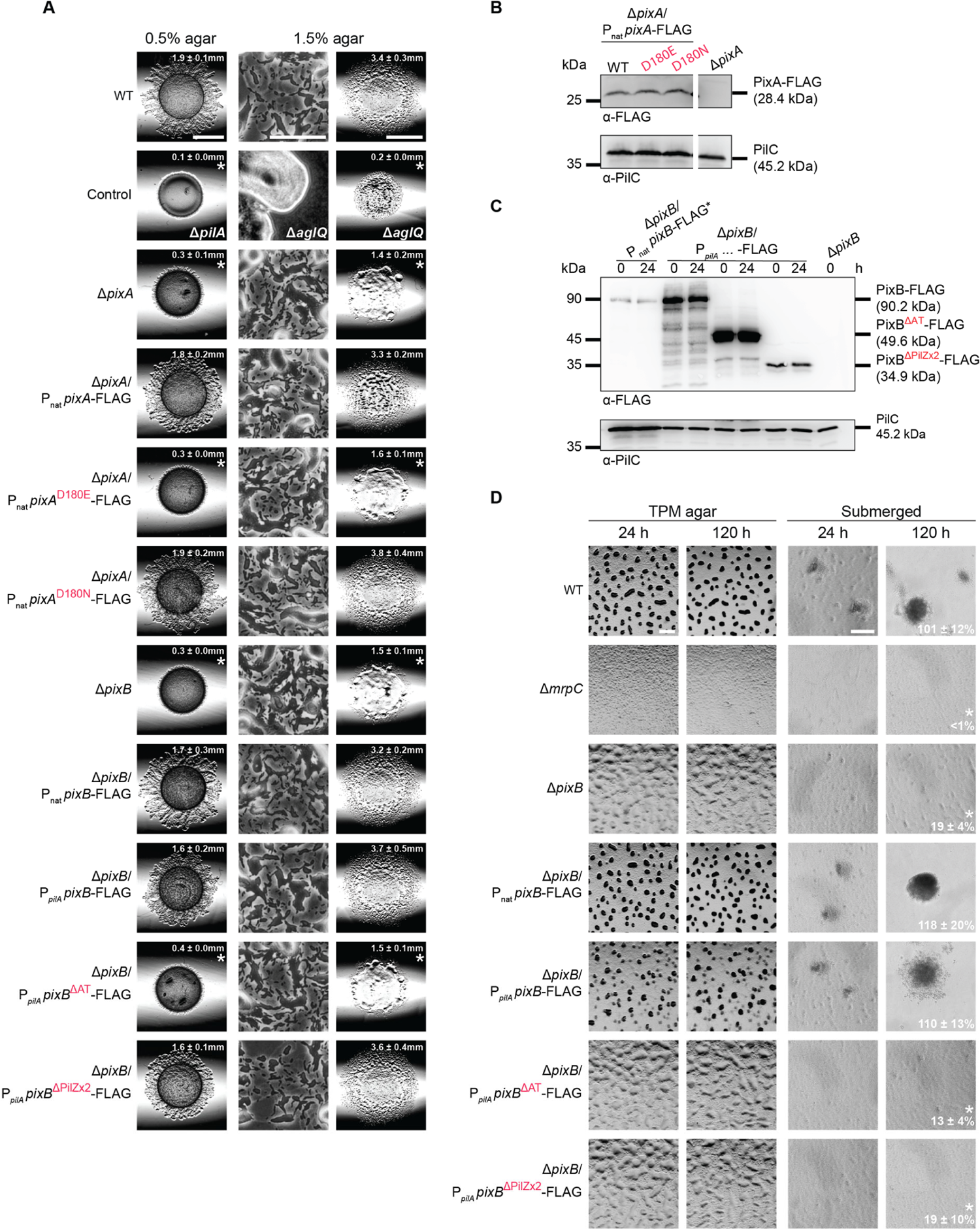
Role of PixA receiver domain and PixB AT domain in regulation of motility and development A. Motility assays of strains expressing mutant variants of PixA and PixB. Motility was analyzed as in Fig. 2. Scale bars, 3 mm (0.5% agar), 100 μm (1.5% agar, left), 3 mm (1.5% agar, right). B, C. Immunoblot detection of PixA-FLAG (B) and PixB-FLAG (C) variants. Samples were prepared and analyzed as in Fig. 7C, D. In B, samples were loaded on the same gel; the gap indicates lanes that were removed for presentation purposes. 10 µg of protein was loaded per lane except for lanes marked * in which 100 µg of protein was loaded and then samples were separated by SDS-PAGE. D. Development assays in strains expressing mutant variants of PixB. Development was analyzed as in Fig. 3. Scale bars, 500 µm (TPM agar), 100 µm (submerged).

To test which domains in PixB are important for function, we ectopically expressed PixB^WT^-FLAG as well as truncated variants that lacked either the AT domain (PixB^ΔAT^-FLAG) or the two PilZ domains (PixB^ΔPilZx2^-FLAG) in the Δ*pixB* mutant. Because the two truncated variants could not be detected by Western blot when expressed from the native *pixB* promoter, all variants were expressed from the stronger *pilA* promoter. The two variants as well as the WT protein accumulated at a higher level than PixB^WT^-FLAG expressed from the native promoter during growth and development (Fig. 8C). Importantly, overexpression of PixB^WT^-FLAG restored both motility and development, while overexpression of PixB^ΔAT^-FLAG restored neither motility nor development (Fig. 8A, D; Fig. S5C, D). Notably, PixB^ΔPilZx2^-FLAG was sufficient to restore motility, but not development (Fig. 8A, D; Fig. S5C, D). Altogether, these observations support that the AT activity is key to PixB function and that c-di-GMP binding to the PilZ domains is important for activating AT activity during development.

## Discussion

Here, we report the identification of two PilZ-domain proteins, PixA and PixB, that are important for regulation of motility in *M. xanthus* and, specifically, important for maintaining a correct cellular reversal frequency during growth. Moreover, PixB is important for completion of the developmental program that results in the formation of spore-filled fruiting bodies.

Similarly to the previously identified PilZ-domain protein PlpA (69), PixA and PixB are not important for motility *per se* but rather are important for regulation of the cellular reversal frequency. Lack of any of the three proteins cause an increase in the reversal frequency suggesting that they all function to inhibit reversals. Importantly, epistasis experiments demonstrated that they act in independent pathways to establish the correct reversal frequency. Moreover, while PixB acts upstream of Frz system, PixA and PlpA act downstream of the Frz system to inhibit reversals.

PixB contains two PilZ domain proteins as well as a C-terminal AT domain. PixB binds c-di-GMP *in vitro*; however, it is not clear which PilZ domain(s) is involved in this binding. By analyzing truncated variants of PixB or variants with single amino acid substitutions in the sequence motifs for c-di-GMP binding, we found that the AT domain was necessary and sufficient for correct reversals during growth. By contrast, the two PilZ domains alone did not complement the reversal defect in the Δ*pixB* mutant. Along the same lines, substitutions in the c-di-GMP binding motifs did not interfere with PixB activity in regulation of the reversal frequency under the conditions tested. These observations support a model whereby the AT activity is key to the effect of PixB on Frz-induced reversals during growth while the two PilZ domains and c-di-GMP binding are not important. How then would the AT domain act to inhibit reversals? Acetyltransferases of the GNAT family can acetylate amino groups of different substrates including small molecules, peptidoglycan, nucleotide-linked monosaccharides and proteins; however, the substrate cannot be determined based on protein sequence (80, 81). Therefore, further studies are required to determine the substrate of PixB.

While the AT domain of PixB was sufficient for correct reversals during growth, all three domains of PixB were important for completion of development. Similarly, PixB variants with single amino acid substitutions in two motifs for c-di-GMP binding did not support development, strongly suggesting that c-di-GMP binding by PixB plays role in fruiting body formation. We previously showed that the level of c-di-GMP increases approximately 10-fold during development (57). We speculate that the increased c-di-GMP level allows binding of c-di-GMP to the two PilZ domains and boost AT activity of PixB during development. Such a mechanism would be similar to that recently described for an AT from *Mycobacterium tuberculosis* that depends on cAMP binding to a cyclic nucleotide binding domain to activate acetyltransferase activity (82). Interestingly, the cellular reversal frequency of WT cells decreases during development (83), thus, it is interesting to speculate that PixB activity is essential for this decrease. In future experiments, it will be of interest to determine the substrate of PixB and to elucidate how the two PilZ domains may regulate PixB activity. Of note, the primary defect in *dmxB* mutants, which lack the DGC responsible for the ∼10-fold increase in c-di-GMP during development, is reduced EPS synthesis (57). The Δ*pixB* mutant accumulates EPS during development at WT levels arguing that the output of DmxB activity is not only channeled through PixB.

PixA and PlpA both act downstream of the Frz system to inhibit reversals and mutants lacking any of the two proteins still accumulate EPS. These observations argue that PixA and PlpA are not involved in the known responses to altered c-di-GMP levels during growth, i.e. altered EPS synthesis and reduced *pilA* transcription. PixA is a response regulator that binds c-di-GMP *in vitro*. Under the conditions tested, c-di-GMP by the PilZ domain was not important for regulation of motility. Moreover, genetic evidence supports that the receiver domain could be phosphorylated and that the non-phosphorylated form represents the active form of the protein. PlpA is a stand-alone PilZ domain protein, which was reported not to bind c-di-GMP *in vitro,* and substitutions in the c-di-GMP binding sequence motifs did not interfere with PlpA activity *in vivo* (69). The Frz-independent pseudo-reversals resulting from lack of PixA and PlpA are characteristic of mutants in which the localization of the small GTPase MglA, the key regulator of motility in *M. xanthus*, in its GTP-bound form is shifted from unipolar to bipolar (46, 49, 77, 84, 85). Consistently, in Δ*plpA* (69) and Δ*pixA* mutants, MglA localization is shifted to bipolar. These observations together with the epistasis experiments strongly argue that PixA and PlpA act in independent pathways at the level of the cell polarity system that establish unipolar T4P and Agl/Glt complex assembly at the leading cell pole; and, in the case of the Agl/Glt complexes, their disassembly at the lagging pole. At the leading cell pole, MglA-GTP stimulates T4P formation and assembly of the Agl/Glt complexes (36, 47, 48) while the MglA GAP MglB at the lagging pole inhibits MglA-GTP accumulation at this pole and, therefore, T4P formation and Agl/Glt complex disassembly (36, 48). Thus, for both motility systems, MglB with its GAP activity at the lagging pole is essential for persistent directional motion without Frz-independent pseudo-reversals. PlpA was reported to localize to the lagging cell pole in an MglB-dependent manner (69) while it is not known whether PixA is polarly localized. In future experiments, it will be interesting to test for interactions between PlpA, PixA and proteins of the polarity module. Genetic evidence suggest that PixA activity could be regulated by phosphorylation of the receiver domain.

Typically, partner proteins of two component systems are encoded by neighboring genes (75); however, in *M. xanthus*, these proteins are are often encoded by orphan genes (86). This is also the case for the *pixA* gene, where there is no neighboring gene encoding a histidine protein kinase. Therefore, it will also be of interest to identify the potential partner kinase of PixA.

Interestingly, *Pseudomonas aeruginosa* as well as *Xanthomonas axonopodis* pv. citri contain a stand-alone PilZ protein that stimulates T4P formation (64, 65). By contrast, in *M. xanthus*, PlpA, PixA and PixB regulate T4P-dependent motility rather than T4P formation. We conclude that PilZ domain proteins can act at different levels to affect T4P-dependent motility.

## Supporting information

All supplemental information

## Acknowledgement

The authors thank Magdalena Anna Świątek-Połatyńska for constructing SA6462. This work was supported by Deutsche Forschungsgemeinschaft (DFG, German Research Council) within the framework of the SFB987 “Microbial Diversity in Environmental Signal Response” and the SPP1879 “Nucleotide second messenger signaling in bacteria” as well as by the Max Planck Society. The funders had no role in study design, data collection and interpretation.

## Data Availability

All the data that support the findings of this study are all included in the manuscript.

## Author contributions

S.K., D.Sk., D.Sc. and L.S.-A. conceptualized the study.

D.Sk., S.K., D.Sc., P.K. and M.P.B. performed bioinformatics studies.

S.K., D.Sk., E.S., P.K. and M.P.B. performed genetic and molecular microbiology experiments.

S.K., D.Sz. and P.K. performed microscopy studies.

D.Sk, D.Sz. and S.K. performed biochemistry experiments.

S.K., D.S. and L.S.-A. wrote the original draft of the manuscript. All authors reviewed and edited the original manuscript.

L.S.-A. acquired funding and provided supervision.

## Materials and methods

### Cultivation of *M. xanthus* and *E. coli*

All *M. xanthus* strains used in this study are derivatives of WT DK1622 (87). In-frame deletions were generated as described (86). All *M. xanthus* strains generated were verified by PCR. *M. xanthus* strains, plasmids and oligonucleotides used are listed in Table 1, Table 2, and Table S1, respectively. *M. xanthus* cells were grown at 32 °C in 1% CTT medium or on 1% CTT/1.5% agar plates at 32  °C with addition of kanamycin (40 µg ml^-1^) or oxytetracycline (10 µg ml^-1^) (88). *E. coli* cells were cultivated in LB (89) liquid media with shaking or on LB 1.5% agar plates at 37°C with addition of kanamycin (40 µg ml^-1^) or tetracycline (10 µg ml^-1^). All plasmids were propagated in *E*. *coli* Top10 (Invitrogen™ life technologies) unless otherwise mentioned.

**Table 1.**
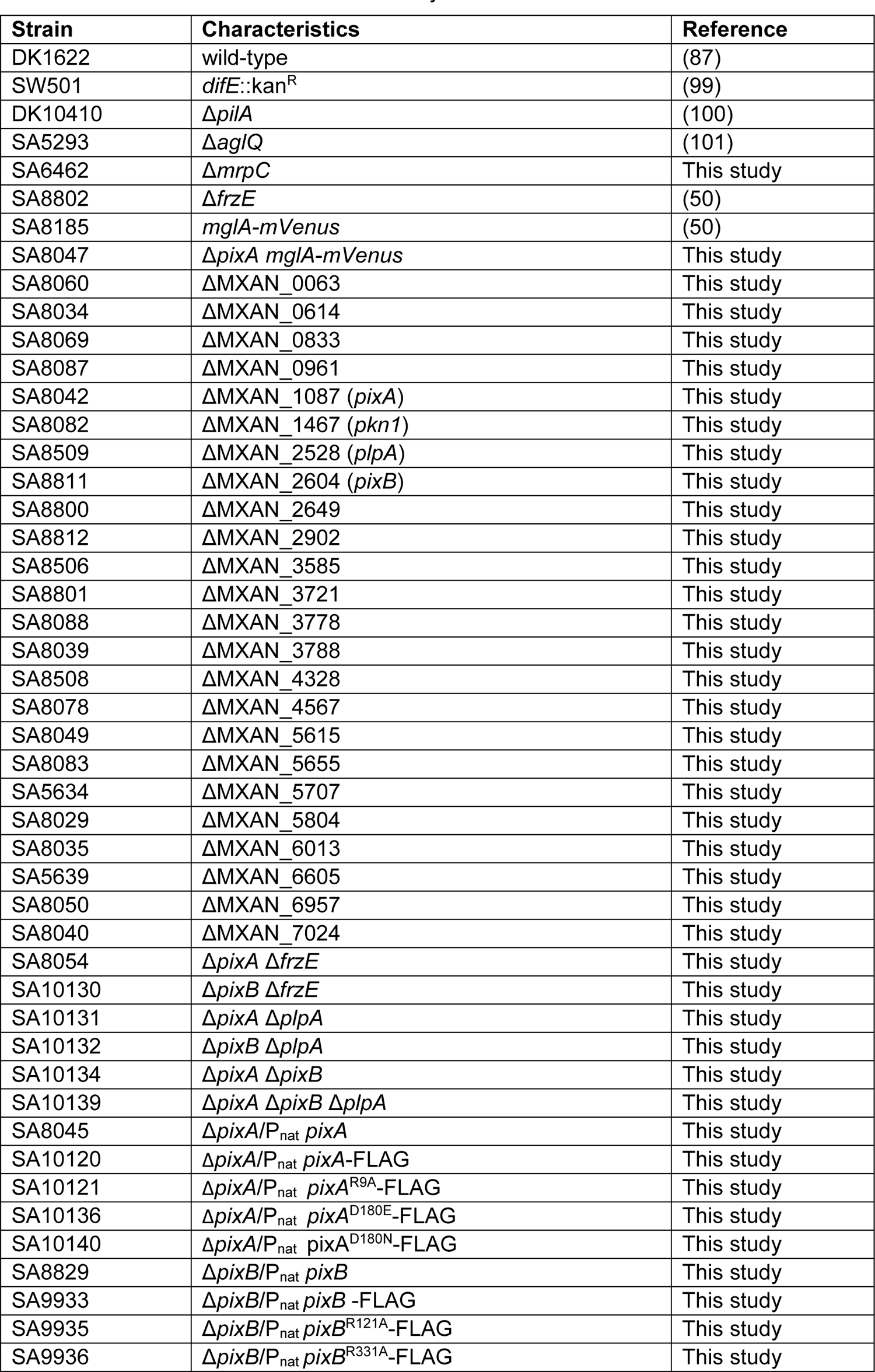
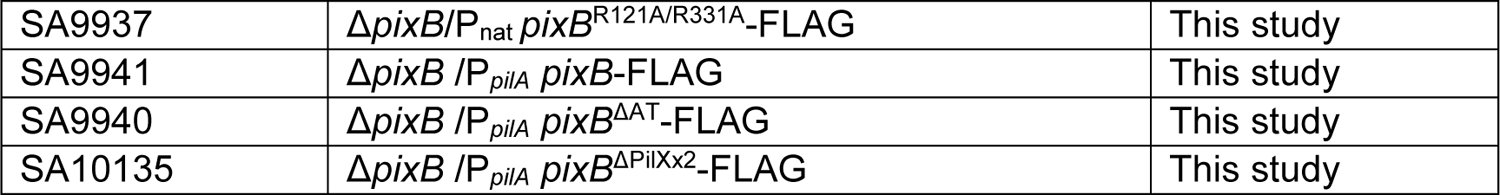
M. xanthus strains used in this study

**Table 2.**
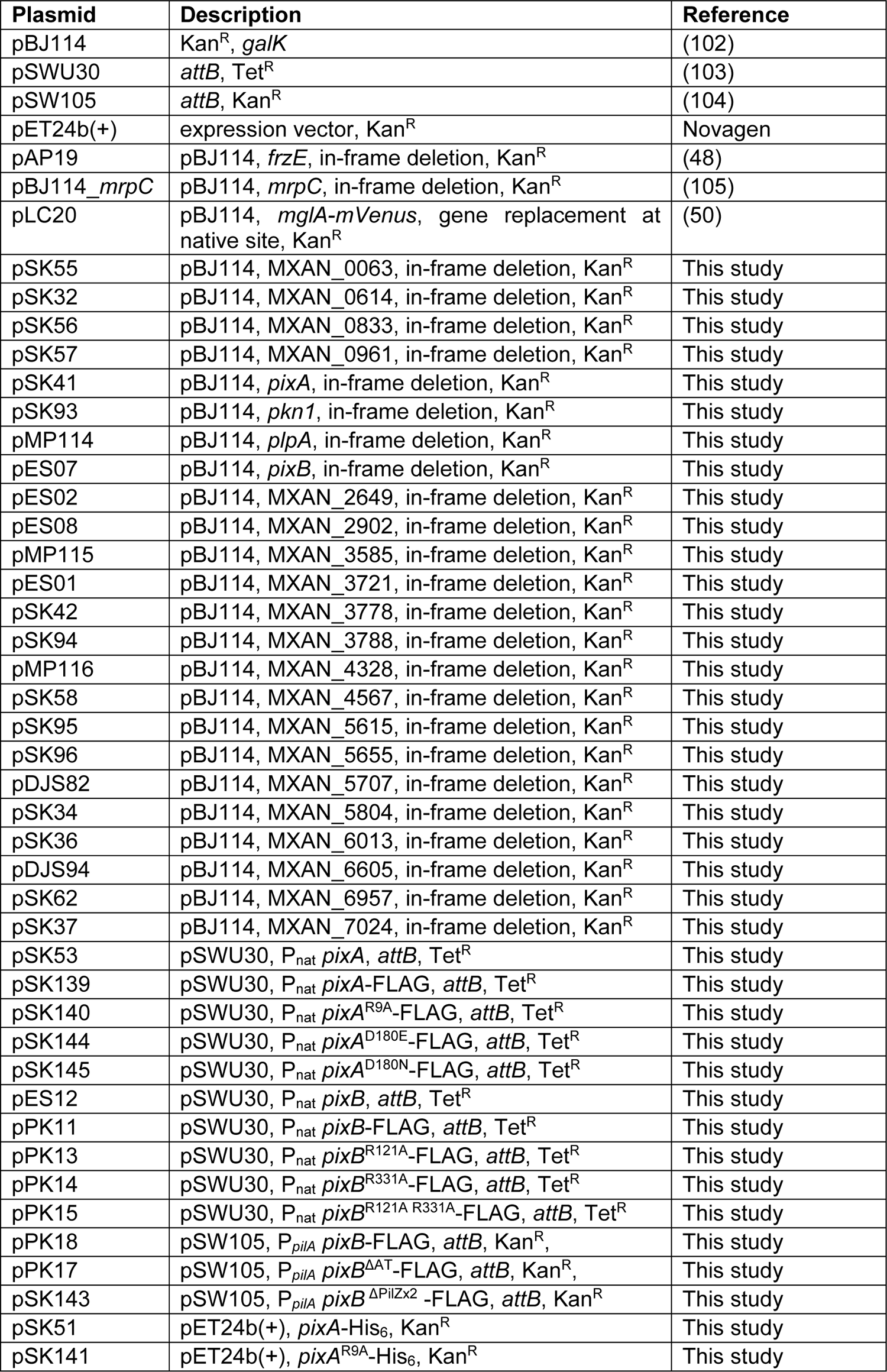
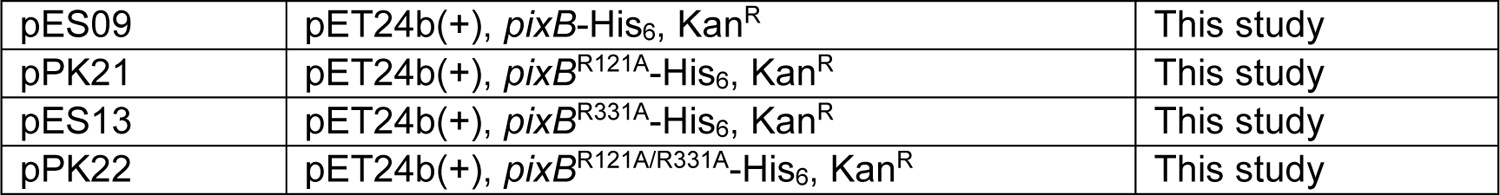
Plasmids used in this study

### Motility assays

Motility assay were done as described with modifications (34). Briefly, exponentially growing *M. xanthus* cells were harvested at 5000 × *g* for 5 min and resuspended in 1% CTT to 7 × 10^9^ cells ml^-1^. From this suspension 5 µl were spotted on 0.5% agar with addition of 1% CTT to final concentration 0.5% for T4P-dependent motility and on 1.5% agar with addition of 1% CTT to final concentration 0.5% for gliding motility. Cells were incubated in dark at 32 °C for 24 h. Colony morphology was imaged using a Leica M205FA Stereomicroscope with a Hamamatsu ORCA-flash V2 Digital CMOS camera and a Leica DMi8 inverted microscope with a Leica DFC280 camera.

### Development. *M*

*xanthus* development assays were performed as described (90) on solid TPM (10 mM Tris-HCl pH 7.6, 1 mM K_2_HPO_4_/KH_2_PO_4_ pH 7.6, 8 mM MgSO_4_) 1.5% agar plates and under MC7 buffer (10 mM MOPS pH 6.8, 1 mM CaCl_2_). Briefly, exponentially growing cells were harvested at 5,000 × *g* for 5 min and resuspended in MC7 buffer to 7 × 10^9^ cells ml^-1^. 20 μl of cell suspension was spotted on TPM agar and 50 μl was added to 350 μl of MC7 buffer in 24-well polystyrene plate (Falcon). At 24 h and 120 h fruiting bodies were imaged with a Leica M205FA stereomicroscope with a Hamamatsu ORCA-flash V2 Digital CMOS camera and a Leica DMi8 inverted microscope with Leica DFC280 camera. To determine sporulation efficiency, cells at 120 h of development were harvested from one of the 24-well polystyrene plate (Falcon). Cells were sonicated 1 min (30% pulse; 50% amplitude with a UP200St sonifier and microtip, Hielscher) to disperse fruiting bodies and then incubated at 55°C for 2 h. Sporulation efficiency was calculated as the number of sonication and heat resistant spores formed after 120 h of development, relatively to WT. Spores were counted in a counting chamber (Depth 0.02 mm, Hawksley).

### Single cell motility assays

Assays were performed as described (50). Briefly, to track individual cells moving by T4P-dependent motility, 5 µl of an exponentially growing cell culture was spotted on 24-well polystyrene plate (Falcon) and incubated at room temperature for 10 min. in the dark. Then cells were covered with 500 µl of 1% methylcellulose in MMC buffer (10 mM MOPS pH 7.6, 4 mM MgSO_4_, 2 mM CaCl_2_) and incubated in the dark at room temperature for 30 min. Cell movement was recorded for 10 min at 20 s intervals. For analysis Metamorph (Molecular Devices) and ImageJ 1.52b (91) were used. For each cell, the distance moved per 20 s interval was determined and the speed per minute calculated; for reversals, the number of reversals per cell per 10 min was determined. Only cells that displayed movement were included in these analyses. To track individual cells moving by gliding, 5 µl of exponentially growing cultures were placed on 1.5% agar plates supplemented with 0.5% CTT, covered by a coverslip and incubated at 32 °C. After 4 h, cells were observed for 10 min at 20 s intervals at RT, and speed per minute as well as the number of reversals per 10 min calculated.

### Trypan Blue dye binding assay

Assay was performed as described (56). Briefly, overnight cultures were grown to a density of 7 × 10^8^ cells ml^-1^ and then were centrifuged at 5,000 × *g* for 5 min and resuspended in MC7 buffer to 7 × 10^9^ cells ml^-1^. 20 μl of cell suspension was spotted on 0.5% agar supplemented with 0.5% CTT or on 1.5% agar plates supplemented with TPM buffer containing Trypan Blue in final concentration of 10 μg/ml. Plates were incubated at 32°C in the darkness for 24 h. Colonies were visualized using a plate scanner.

### Microscopy and analysis of fluorescence microscopy images

Epifluorescence microscopy was performed as described (50). Briefly, exponentially growing cells were placed on a thin 1.5% agar pad buffered with TPM buffer on a glass slide and immediately covered with a coverslip. After 30 min at 32 °C, cells were visualized using a Leica DMi8 inverted microscope and phase contrast and fluorescence images acquired using a Leica DFC280 camera. Cells in phase contrast images were automatically detected using Oufti48.

Fluorescence signals in segmented cells were identified and analyzed using a custom-made Matlab v2016b (MathWorks) script as described in (50). The script divides a cell into three regions: polar region 1, polar region 2 and the cytoplasmic region. The polar regions are defined as the parts of a cell within a distance of 10 pixels, corresponding to 0.64 µm, from a tip of a cell. The cytoplasmic region includes all pixels of the cell with the exception of the polar regions. A polar cluster was identified when three or more connected pixels within a polar region had a fluorescence signal higher than a cell-specific threshold signal of two standard deviations above the average fluorescence signal in the cytoplasmic region. The fluorescence of a polar cluster was defined as the sum of the fluorescence signal of all connected pixels that exceeded the threshold value in that polar region. The cytoplasmic signal was defined as the sum of the fluorescence signal of all pixels excluding the polar clusters. For each cell with polar cluster(s), an asymmetry index (ω) was calculated as

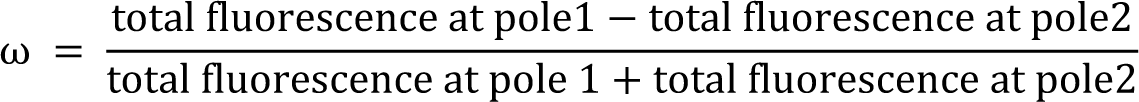

By definition, pole 1 is the pole with the highest fluorescence. ω varies between 0 (bipolar symmetric localization) and 1 (unipolar localization). The localization patterns were binned from the ω values as follows: unipolar (ω > 0.9), bipolar asymmetric (0.9 > ω > 0.2) and bipolar symmetric (ω < 0.2). Diffuse localization was determined when no polar signal was detected. For time-lapse epifluorescence microscopy, cells were prepared as described and recorded for 15 min with images captured every 30 s. Data were processed with Metamorph 7.5 and ImageJ 1.52b.

### Immunoblot analysis

Protein concentration was measured using a Bradford assay (BioRad). Rabbit polyclonal α-PilC (1:5000 dilution) (Bulyha et al., 2009) and α-FLAG (Rockland, 1:1500 dilution) antibodies were used together with horseradish-conjugated goat α-rabbit immunoglobulin G (Sigma-Aldrich) as secondary antibody. Immunoblots were performed as described (89). Blots were developed using Luminata crescendo Western HRP Substrate (Millipore) and visualized with a LAS-4000 luminescent image analyzer (Fujifilm).

### Protein purification

To purify PixA-His_6_, PixA^R9A^-His_6_, PixB-His_6_, PixB^R121A^-His_6_ and PixB^R331A^-His_6_, PixB^R121A/R331^-His_6_ and PixB-His_6_, *E*. *coli* Rosseta 2 (DE3)/pLysS strain (Novagen) was transformed with pSK51, pSK141, pES09, pPK21, pES13 and pPK22 respectively. Cultures were grown in 1 L of LB with addition of chloramphenicol and kanamycin at 37 °C to an OD_600_ of 0.5-0.7. Protein expression was induced by addition of Isopropyl β-d-1-thiogalactopyranoside (IPTG) to a final concentration of 0.5 mM for 3 h at 37°C. Cells were harvested by centrifugation at 3,800 × *g* for 10 min at 4 °C and resuspended in lysis buffer (50 mM NaH_2_PO_4_, 300 mM NaCl, 50 mM imidazole, 5% glycerol and Complete Protease Inhibitor Cocktail Tablet (Roche), pH 8.0). Next, cells were disrupted with French press and centrifuged at 48,000 × *g*, 4°C for 40 min. Cleared cell lysate was filter with 0.45 µm sterile filter (Millipore Merck, Schwalbach) and loaded onto a 5 ml HiTrap Chelating HP column (GE Healthcare) preloaded with NiSO_4_ as described by the manufacturer and pre-equilibrated in wash buffer (50 mM NaH_2_PO_4_, 300 mM NaCl, 50 mM imidazole, 5% glycerol, pH 8.0). The column was washed with 20 column volumes of column wash buffer. Proteins were eluted with elution buffer A (50 mM NaH_2_PO_4_, 300 mM NaCl, 500 mM imidazole, 5% glycerol, pH 8.0) using a linear imidazole gradient from 50 to 500 mM. Fractions containing purified His_6_-tagged proteins were combined and loaded onto a HiLoad 16/600 Superdex 75 pg (GE Healthcare) gel filtration column that was equilibrated with buffer (50 mM NaH_2_PO_4_, 300 mM NaCl, 5% glycerol, pH 6.5). Fractions containing His_6_-tagged proteins were pooled, frozen in liquid nitrogen and stored at −80 °C.

### *In vitro* c-di-GMP binding assay

c-di-GMP binding was determined using a DRaCALA assay with ^32^P-labeled c-di-GMP as described (92, 93). ^32^P-labeled c-di-GMP was prepared by incubating 10 µM His_6_-DgcA with 1 mM GTP/[α-^32^P]-GTP (0.1 μCi μl^−1^) in reaction buffer (total volume of 200 µl) overnight at 30°C. The reaction mixture was then incubated with 5 units of Antarctic Phosphatase (NEB) for 1 h at 22 °C to hydrolyze unreacted GTP. The reaction was stopped by incubation for 10 min at 95 °C. The reaction was centrifuged (10 min., 20,000 × *g*, 20 °C) and the supernatant used for the binding assay. ^32^P-labeled c-di-GMP was mixed with 20 µM protein and incubated for 10 min at RT in binding buffer (10 mM Tris, pH 8.0, 100 mM NaCl, 5 mM MgCl_2_). 15 µl of this reaction mixture was transferred to a nitrocellulose filter, allowed to dry and imaged using a STORM 840 Scanner (Amersham Biosciences). For competition experiments, 0.4 mM unlabelled c-di-GMP (BioLog) or GTP (Sigma-Aldrich) was added.

### Bioinformatic analysis

The KEGG SSDB (Sequence Similarity Database) (94) database was used to identify homologs of the PilZ domain proteins in other fully sequenced Myxococcales genomes using a reciprocal best BlastP hit method. Protein domains were identified using Pfam v33.1 (pfam.xfam.org) (95) and MyHits Motif scan (https://myhits.isb-sib.ch/cgi-bin/motif_scan) (96). Multiple Alignment using Fast Fourier Transform from EMBL-EBI (97) was used to align protein sequences. Sequence identity/similarity was calculated using EMBOSS Needle software (98) (pairwise sequence alignment).

