## Supplementary material for "Three PilZ domain proteins, PlpA, PixA and PixB, have distinct functions in regulation of motility and development in *Myxococcus xanthus*": All supplemental information

This file contains:

- Supplementary Figures 1-5
- Supplementary Materials and Methods
- Supplementary Table 1
- Supplementary References

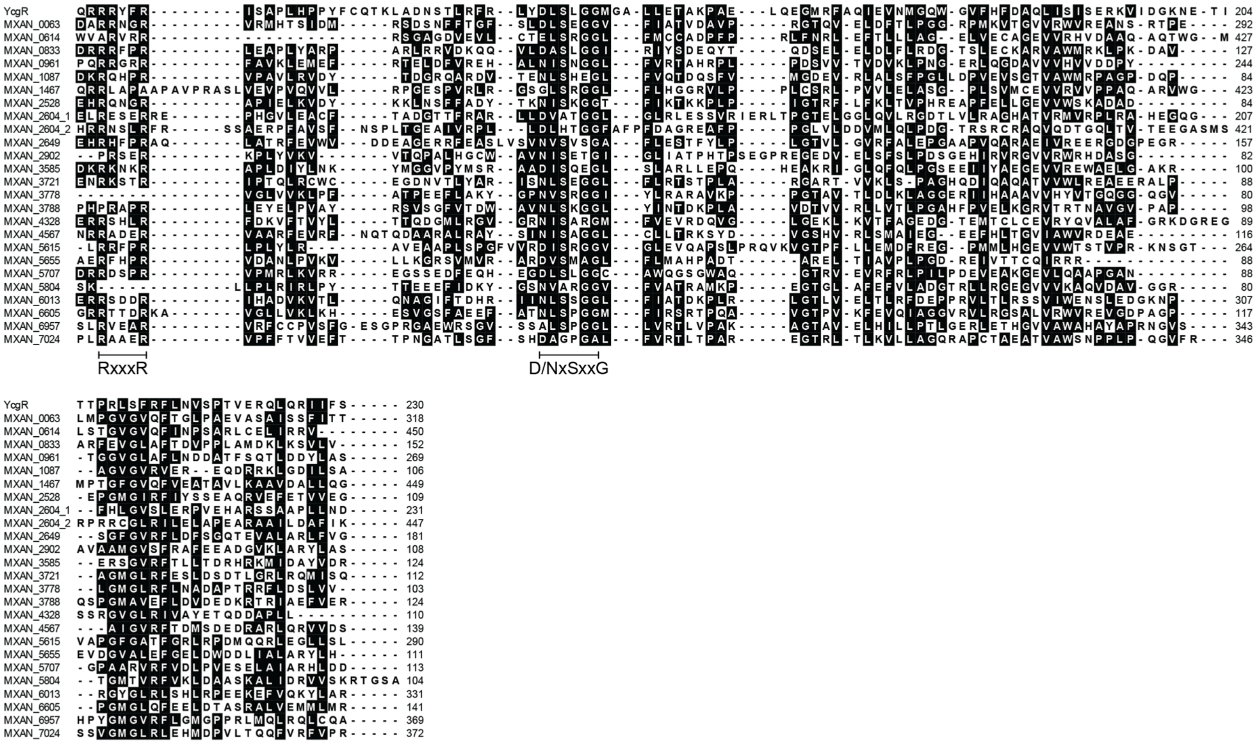


**Figure S1. Sequence alignment of PilZ domains of *M. xanthus*.**
Amino acid sequence of the *M. xanthus* PilZ domain were aligned with YcgR from *E. coli* (P76010) (1), which was also used as a reference to identify the residues important for c-di-GMP binding. The consensus sequence motifs for c-di-GMP binding are shown below. Identical and similar amino acids are marked on black.

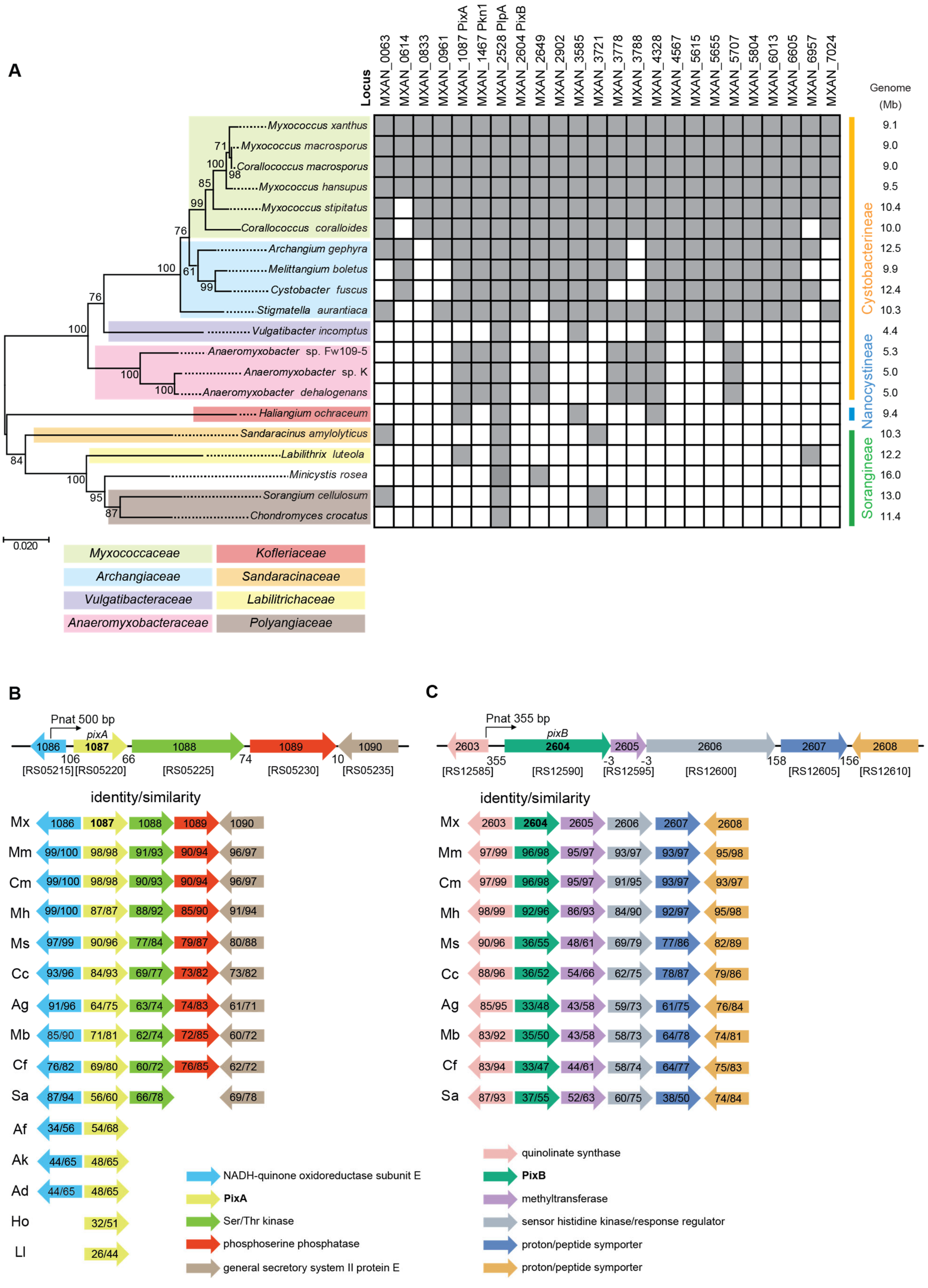
A
Locus
MXAN_0063
MXAN_0614
MXAN_0833
MXAN_0961
MXAN_1087 PixA
MXAN_1467 Pkn1
MXAN_2528 PlpA
MXAN_2604 PixB
MXAN_2649
MXAN_2902
MXAN_3585
MXAN_3721
MXAN_3778
MXAN_3788
MXAN_4328
MXAN_4567
MXAN_5615
MXAN_5655
MXAN_5707
MXAN_5804
MXAN_6013
MXAN_6605
MXAN_6957
MXAN_7024
Genome (Mb)
Myxococcus xanthus 9.1
Myxococcus macrosporus 9.0
Corallococcus macrosporus 9.0
Myxococcus hansupus 9.5
Myxococcus stipitatus 10.4
Corallococcus coralloides 10.0
Archangium gephyra 12.5
Melittangium boletus 9.9
Cystobacter fuscus 12.4
Stigmatella aurantiaca 10.3
Vulgatibacter incomptus 4.4
Anaeromyxobacter sp. Fw109-5 5.3
Anaeromyxobacter sp. K 5.0
Anaeromyxobacter dehalogenans 5.0
Haliangium ochraceum 9.4
Sandaracinus amylolyticus 10.3
Labilithrix luteola 12.2
Minicystis rosea 16.0
Sorangium cellulosum 13.0
Chondromyces crocatus 11.4
71
100
98
85
99
76
100
61
99
76
100
100
100
84
100
95
87
Cystobacterineae
Nanocystineae
Sorangineae
0.020
Myxococcaceae
Archangiaceae
Vulgatibacteraceae
Anaeromyxobacteraceae
Kofleriaceae
Sandaracinaceae
Labilitrichaceae
Polyangiaceae
B
Pnat 500 bp
pixA
1086
1087
1088
1089
1090
106
66
74
10
[RS05215]
[RS05220]
[RS05225]
[RS05230]
[RS05235]
identity/similarity
Mx 1086 1087 1088 1089 1090
Mm 99/100 98/98 91/93 90/94 96/97
Cm 99/100 98/98 90/93 90/94 96/97
Mh 99/100 87/87 88/92 85/90 91/94
Ms 97/99 90/96 77/84 79/87 80/88
Cc 93/96 84/93 69/77 73/82 73/82
Ag 91/96 64/75 63/74 74/83 61/71
Mb 85/90 71/81 62/74 72/85 60/72
Cf 76/82 69/80 60/72 76/85 62/72
Sa 87/94 56/60 66/78 69/78
Af 34/56 54/68
Ak 44/65 48/65
Ad 44/65 48/65
Ho 32/51
Ll 26/44
NADH-quinone oxidoreductase subunit E
PixA
Ser/Thr kinase
phosphoserine phosphatase
general secretory system II protein E
C
Pnat 355 bp
pixB
2603
2604
2605
2606
2607
2608
355
-3
-3
158
156
[RS12585]
[RS12590]
[RS12595]
[RS12600]
[RS12605]
[RS12610]
identity/similarity
Mx 2603 2604 2605 2606 2607 2608
Mm 97/99 96/98 95/97 93/97 93/97 95/98
Cm 97/99 96/98 95/97 91/95 93/97 93/97
Mh 98/99 92/96 86/93 84/90 92/97 95/98
Ms 90/96 36/55 48/61 69/79 77/86 82/89
Cc 88/96 36/52 54/66 62/75 78/87 79/86
Ag 85/95 33/48 43/58 59/73 61/75 76/84
Mb 83/92 35/50 43/58 58/73 64/78 74/81
Cf 83/94 33/47 44/61 58/74 64/77 75/83
Sa 87/93 37/55 52/63 60/75 38/50 74/84
quinolinate synthase
PixB
methyltransferase
sensor histidine kinase/response regulator
proton/peptide symporter
proton/peptide symporter

**Figure S2. Conservation of PilZ domain-containing proteins of *M. xanthus* in other myxobacteria.**

A. Taxonomic distribution of *pilZ* genes in Myxococcales with fully sequenced genomes. A reciprocal best BlastP hit method was used to identify *pilZ* gene orthologs in myxobacteria. MXAN locus tags and gene names are indicated. Grey indicates conserved genes while white indicates that no homolog was identified. Left, 16S rRNA tree of Myxococcales with fully sequenced genomes was prepared based as in (2). Bootstrap values (500 replicates) are shown next to the branches (3). Family and suborder classification are indicated. Genome size is indicated on the right.

B, C. Locus organization of *pixA* (B) and *pixB* (C). Top panel: arrows indicate direction of transcription. Distance between start and stop codon of neighboring genes is indicated. The kinked arrows indicate the fragments referred to as the native promoters ($P_{nat}$), which were used to express the genes ectopically from the *attB* site. Bottom panel: conservation of the loci in fully sequenced Myxococcales genomes. Numbers in the arrows indicate % identity/similarity between proteins from *M. xanthus* and their homologs. Species abbreviations can be taken from (A).

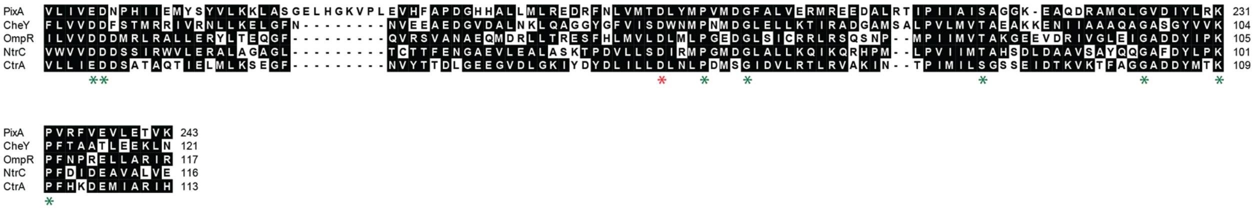

**Figure S3. Sequence alignment of the receiver domain of PixA with receiver domains of NtrC (P0AFB8), OmpR (P0AA16), CtrA (B8H358) and CheY (P0AE67).**
Identical and similar residues are marked on black. Residues important for phosphorylation (4) are marked with green asterisks and the phosphorylatable Asp residue is marked with a red asterisk.

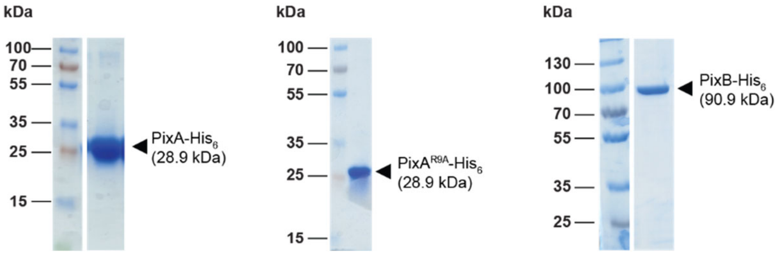


**Figure S4. Purification of His6-tagged PixA and PixB variants.**
Purified PixA-His$_6$ and PixB-His$_6$ protein variants were separated by SDS-PAGE and stained with Coomassie blue. Calculated molecular mass of the different proteins is indicated. Molecular size markers are indicated on the left.

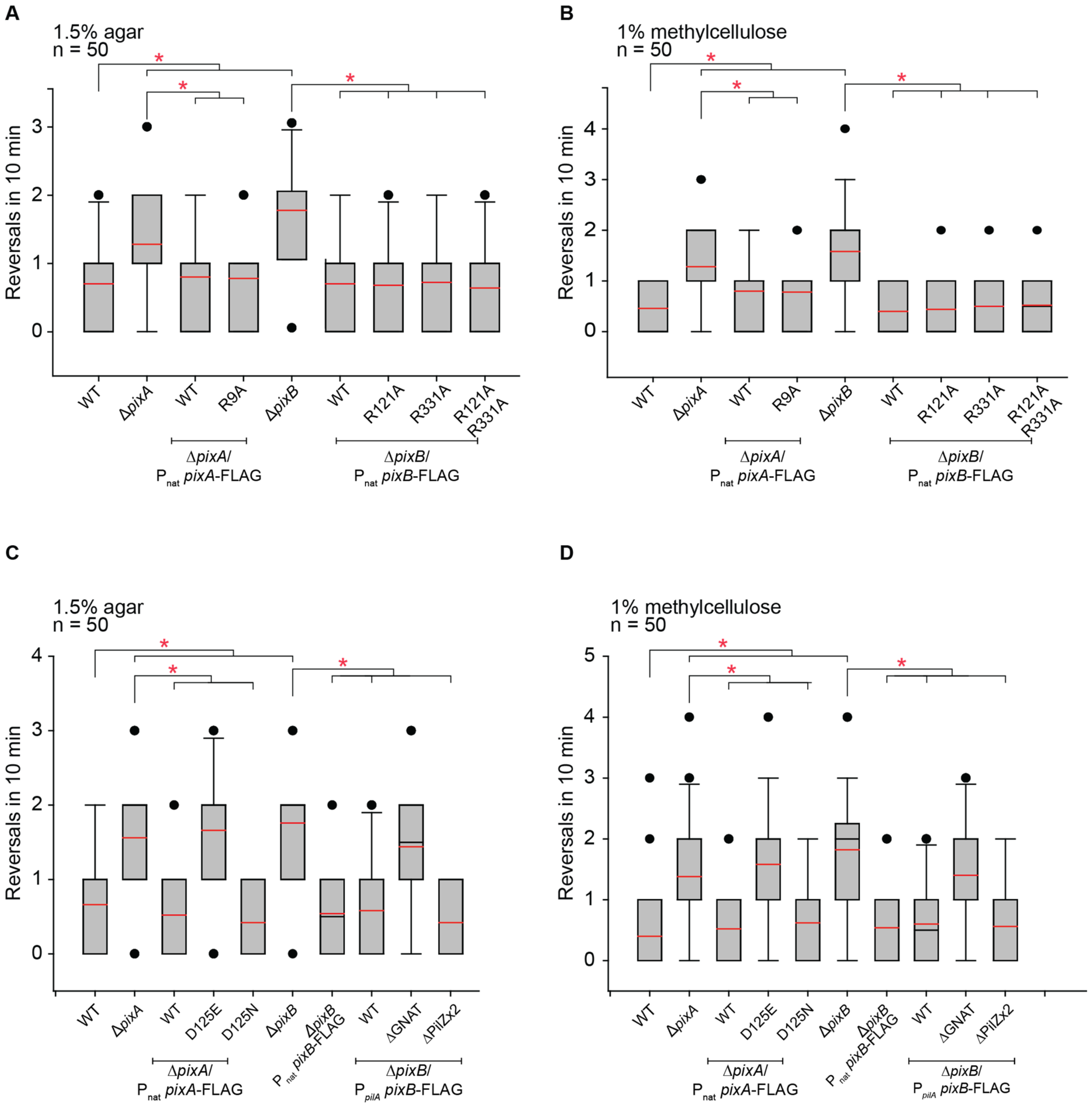


**Figure S5. Single cell reversal frequency of strains of indicated genotypes.**
A-D. Reversals of indicated strains on 1.5% agar supplemented with 0.5% CTT (A, C) or in 1% methylcellulose (B, D). Boxplots are as in Fig. 4D, F. * $P < 0.05$ in Mann-Whitney Rank Sum Test, n = 50 cells.

**Supplementary Materials and Methods**

Plasmid construction:

The plasmids pSK32, pSK34, pSK36, pSK37, pSK41, pSK42, pSK55, pSK56, pSK57, pSK58, pSK62, pSK93, pSK94, pSK95, pSK96, pMP114, pMP115, pMP116, pES01, pES02, pES07, pES08, pDJS82, pDJS94 were generated for the construction of the MXAN_0614, MXAN_5804, MXAN_6013, MXAN_7024, MXAN_1087 (*pixA*), MXAN_3778, MXAN_0063, MXAN_0833, MXAN_0961, MXAN_4567, MXAN_6957, MXAN_1467 (*pkn1*), MXAN_3788, MXAN_5615, MXAN_5655, MXAN_2528 (*plpA*), MXAN_3585, MXAN_4328, MXAN_3721, MXAN_2649, MXAN_2604 (*pixB*), MXAN_2902, MXAN_5707 and MXAN_6605 in-frame deletion mutants respectively. The upstream and downstream regions of the genes were amplified using primer pairs SK82-SK61 and SK62-SK63, SK036-SK037 and SK038-SK039, SK72-73 and SK74-75, SK76-SK77 and SK78-SK79, SK64-SK65 and SK66-SK67, SK68-SK69 and SK70-SK71, SK156- SK157 and SK158- SK159, SK144-SK145 and SK146-SK147, SK160-SK161 and SK162-SK163, SK164-SK165 and SK166-SK167, SK148-SK149 and SK150-SK151, SK253-SK254 and SK255-SK256, SK245-SK246 and SK247-SK248, SK152-SK261 and SK252-SK155, SK237-SK238 and SK239-SK240, 2528_A-2528_B and 2528_C-2528_D, 3585_A-3585_B and 3585_C-3585_D, 4328_A-4328_B and 4328_C-4328_D, 3721_A-3721_B and 3721_C-3721_D, 2649_A-2649_B and 2649_C-2649_D, 2604_A-2604_B and 2604_C-2604_D, 2902_A-2902_B and2902_C-2902_D, 5707_A-5707_B and 5707_C-5707_D, 6605_A-6605_B and 6605_C-6605_D. The AB and CD DNA fragments were fused by overlap PCR reaction. Resulting AD fragments were cloned into KpnI/XbaI or EcoRI/XbaI site of pBJ114.

Plasmid **pSK53** was used to complement Δ*pixA* mutant with WT version of *pixA* under the native promoter. To amplify *pixA* with the native promoter primers SK128-SK129 were used. The product was cloned into HindIII/XbaI sites of pSWU30.

Plasmid **pSK139** was used to complement Δ*pixA* mutant with full-length version of *pixA* fused to FLAG-tag under the native promoter. To amplify *pixA*-FLAG with the native promoter primers SK128 - SK384 were used. The product was cloned into HindIII/XbaI sites of pSWU30.

Plasmid **pSK140** was used to complement Δ*pixA* mutant with full-length version of *pixA*$^{R9A}$ fused to FLAG-tag under the native promoter. To amplify *pixA*$^{R9A}$-FLAG with the native promoter primers SK128 - SK385 and SK386 - SK384 were used. Obtained fragments were fused during second round of PCR amplification using primers SK128 - SK384. The product was cloned into HindIII/XbaI sites of pSWU30.

Plasmid **pSK144** was used to complement Δ*pixA* mutant with full-length version of *pixA*$^{D180E}$ fused to FLAG-tag under the native promoter. To amplify *pixA*$^{D180E}$-FLAG with the native promoter primers SK64 – SK190 and SK189 - SK384 were used. Obtained fragments were fused during second round of PCR amplification using primers SK64 - SK384. The product was cloned into HindIII/EcoRI sites of pSWU30.

Plasmid **pSK145** was used to complement Δ*pixA* mutant with full-length version of *pixA*$^{D180N}$ fused to FLAG-tag under the native promoter. To amplify *pixA*$^{D180N}$-FLAG with the native promoter primers SK64 – SK192 and SK191 - SK384 were used. Obtained fragments were fused during second round of PCR amplification using primers SK64 - SK384. The product was cloned into HindIII/EcoRI sites of pSWU30.

Plasmid **pES12** was used to complement Δ*pixB* mutant with WT version of *pixB* under the native promoter. To amplify *pixB* with the native promoter primers Pnat 2604 XbaI fw - 2604 KpnI rev were used. The product was cloned into XbaI/KpnI sites of pSWU30.

Plasmid **pPK11** was used to complement Δ*pixB* mutant with full-length version of *pixB* fused to FLAG-tag under the native promoter. To amplify *pixB*-FLAG with the native promoter primers Pnat 2604 XbaI fw - 2604 FLAG Rw were used. The product was cloned into HindIII/XbaI sites of pSWU30.

Plasmid **pPK13** was used to complement Δ*pixB* mutant with full-length version of *pixB*$^{R121A}$ fused to FLAG-tag under the native promoter. To amplify *pixB*$^{R121A}$-FLAG with the native promoter primers Pnat 2604 XbaI fw - R121A PilZ1 and R121A PilZ1 2604 fw - 2604 FLAG Rw were used. Obtained fragments were fused during second round of PCR amplification using primers Pnat 2604 XbaI fw - 2604 FLAG Rw. The product was cloned into HindIII/XbaI sites of pSWU30.

Plasmid **pPK14** was used to complement Δ*pixB* mutant with full-length version of *pixB*$^{R331A}$ fused to FLAG-tag under the native promoter. To amplify *pixB*$^{R331A}$-FLAG with the native promoter primers Pnat 2604 XbaI fw - R331A 2604 rev and R331A 2604 fw - 2604 FLAG Rw were used. Obtained fragments were fused during second round of PCR amplification using primers Pnat 2604 XbaI fw - 2604 FLAG Rw. The product was cloned into HindIII/XbaI sites of pSWU30.

Plasmid **pPK15** was used to complement Δ*pixB* mutant with full-length version of *pixB*$^{R121A/R331A}$ fused to FLAG-tag under the native promoter. To amplify MXAN_2604$^{R331A}$-FLAG with the native promoter primers Pnat 2604 XbaI fw - R331A 2604 rev and R331A 2604 fw - 2604 FLAG Rw were used. As a template **pPK13** was used. Obtained fragments were fused during second round of PCR amplification using primers Pnat 2604 XbaI fw - 2604 FLAG Rw. The product was cloned into HindIII/XbaI sites of pSWU30.

Plasmid **pPK18** was used to complement Δ*pixB* mutant with full-length version of *pixB* fused to FLAG-tag under the *pilA* promoter. To amplify *pixB*-FLAG primers 2604 XbaI fw – 2604 FLAG Rw were used. The product was cloned into HindIII/XbaI sites of pSW105.

Plasmid **pPK17** was used to complement Δ*pixB* mutant with the version of *pixB* with deleted AT domain and fused to FLAG-tag under the *pilA* promoter. To amplify *pixB*$^{\Delta AT}$-FLAG primers 2604 XbaI fw - 2604 ΔGNAT-FLAG (451) rev were used. The product was cloned into HindIII/XbaI sites of pSW105.

Plasmid **pSK143** was used to complement Δ*pixB* mutant with the version of *pixB* with deleted PilZ domains and fused to FLAG-tag under the *pilA* promoter. To amplify *pixB*$^{\Delta PilZx24}$-FLAG primers SK387 - 2604 FLAG Rw were used. The product was cloned into HindIII/XbaI sites of pSW105.

Plasmid **pSK51** was used to purify PixA-His$_6$ from *E.coli*. To amplify *pixA* primers SK122 and SK123 were used. The product was cloned into NdeI/XhoI sites of pET24b(+).

Plasmid **pSK141** was used to purify PixA$^{R9A}$-His$_6$ from *E.coli*. To amplify *pixA*$^{R9A}$ primers SK122 and SK123 were used on **pSK140** as a template. The product was cloned into NdeI/XhoI sites of pET24b(+).

Plasmid **pES09** was used to purify PixB-His$_6$ from *E.coli*. To amplify *pixB* primers 2604 NdeI fw – 2604 HindIII –stop rev were used. The product was cloned into NdeI/HindIII sites of pET24b(+).

Plasmid **pPK21** was used to purify PixB$^{R121A}$-His6 from E.coli. To amplify PixB$^{R121A}$ primers 2604 NdeI fw - R121A PilZ1 2604 rev and R121A PilZ1 2604 fw - 2604 HindIII –stop rev were used. Obtained fragments were fused during second round of PCR amplification using primers 2604 NdeI fw - 2604 HindIII –stop rev. The product was cloned into NdeI/HindIII sites of pET24b(+).

Plasmid **pES13** was used to purify PixB$^{R331A}$-His$_6$ from *E.coli*. To amplify *pixB*$^{R331A}$ primers 2604 NdeI fw - R331A 2604 rev and R331A 2604 fw - 2604 HindIII –stop rev were used. Obtained fragments were fused during second round of PCR amplification using primers 2604 NdeI fw – 2604 HindIII –stop rev. The product was cloned into NdeI/HindIII sites of pET24b(+).

Plasmid **pPK22** was used to purify PixB$^{R121A/R331A}$-His6 from E.coli. To amplify PixB$^{R121A/R331A}$ primers 2604 NdeI fw - R121A PilZ1 2604 rev and R121A PilZ1 2604 fw - 2604 HindIII –stop rev were used on **pPK14** DNA. Obtained fragments were fused during second round of PCR amplification using primers 2604 NdeI fw - 2604 HindIII –stop rev. The product was cloned into NdeI/HindIII sites of pET24b(+).

**Table S1. Primers used in this study**

| Name | Sequence (5‘-3‘) | Description |
|---|---|---|
| SK156_A | GCGCGGTACCGACCGCACCTTCGCGGACGT | In-frame deletion of MXAN_0063 |
| SK157_B | TCAGTCCGGGAAGAACATTCCCCCGGACGGCCCCAC | |
| SK158_C | GTGGGGCCGTCCGGGGGAATGTTCTTCCCGGACTG | |
| SK159_D | GCGCAAGCTTGCTTCACGTCCTCCAGGCCG | |
| SK201_E | GGCATCCTCCTCAATACCCG | |
| SK202_F | GTGCAGTTCAGGTGGTGGAT | |
| SK203_G | AAGCAGGTTCGGAGATGCAG | |
| SK204_H | CGCTGAGGTCCAGTGAGAAA | |
| SK82_A | GCGCGAATTCCCGCGTGCGCGGAGACGC | In-frame deletion of MXAN_0614 |
| SK61_B | CACCTCCGCCCGGGCGCGCATCACCCC | |
| SK62_C | CGCGCCCGGGCGGAGGTGGTGCTGTGA | |
| SK63_D | GCGCTCTAGACGAACCACCTGCCCGGCC | |
| SK085_E | ATCGAGCGGCGCACCTGC | |
| SK086_F | GGCACGTCTACGTGAAGATGGACG | |
| SK087_G | GGCGTGCCCATGTCGCAC | |
| SK088_H | CCGCCGCCGTTTCATCCT | |
| SK144_A | GCGCGAATTCAGGCCGGTGACGTTGAGG | In-frame deletion of MXAN_0833 |
| SK145_B | CTATGAGTCGGCCGGGCCAGCCAGCCCGTGGCACA | |
| SK146_C | GTGTGCCACGGGCTGGCTGGCCCGGCCGACTCATA | |
| SK147_D | GCGCTCTAGACTCAGAGGGCGGGGCTCA | |
| SK176_E | ACCTCTCGTTGGGGGATGAC | |
| SK177_F | GCACGGGATGAAATGGTCG | |
| SK178_G | CTGGCTGGGTTCTGGGATTT | |
| SK179_H | GAGCACCGACTTGAGCTTGT | |
| SK160_A | GCGCGAATTCCCACGCGCGGGTCCGTCTCG | In-frame deletion of MXAN_0961 |
| SK161_B | TCAGCCCACGCCACCCGCAAGACCGAAGACGGGCA | |
| SK162_C | GTGCCCGTCTTCGGTCTTGCGGGTGGCGTGGGCTG | |
| SK163_D | GCGCTCTAGATGGAGGGCGACCCGGGCATGGTGC | |
| SK180_E | TCCGGGGCGAGCGGGTTGGTGCC | |
| SK181_F | TCGGGGACGCCGGGCACACGCA | |
| SK182_G | TGGACCACCACCGCGTCGCC | |
| SK183_H | CCGCGGCCTCCGTGCCTCAA | |
| SK64_A | GCGCGAATTCTCTCTCGCAAGGTGAACT | In-frame deletion of MXAN_1087 (*pixA*) |
| SK65_B | GCGCAAGAGCTCTGGACCCGGGTTCAT | |
| SK66_C | GGTCCAGAGCTCTTGCGCATCAAGTAG | |
| SK67_D | GCGCTCTAGAGCCTCGTCCAGCAGCAGC | |
| SK102_E | CTGGGTCAGGCTTTCGTGGTGAT | |
| SK103_F | CATGGCCGAGCGCGAGCT | |
| SK104_G | GGTCCGGCTCGCACTTTCTT | |
| SK105_H | GGACCTCCAGGGGGACCTTT | |
| SK253_A | GCCGGGTACCAAGAAGGGCATCGAGTAGCGCACC | In-frame deletion of MXAN_1467 (*pkn1*) |
| SK254_B | TCAAGGTGACCGGGCCCTGCTGCTCACCTCGGGCAT | |

| | | |
|---|---|---|
| SK255_C | ATGCCCGAGGTGAGCAGCAGGGCCCGGTCACCTTG | |
| SK256_D | GCCG<u>TCTAGA</u>GCCGGCGATGGAGGTGACCTT | |
| SK257_E | TGCCCGTGCGGTAATGGGAC | |
| SK258_F | GCAAGCGCGGCCTCGATAAG | |
| SK259_G | AACGCGCCCTTCGTCAAGGT | |
| SK260_H | CGCGATGGAGGCGTATCGGT | |
| 2528_A | ATAT<u>GGTACC</u>CTCCAAGCGGAACCAGCCGA | In-frame deletion of MXAN_2528 (*plpA*) |
| 2528_B | CGGCTTGTTCTCTGGACCTGTCTTCTG | |
| 2528_C | GGTCCAGAGAACAAGCCGCTGCACTCG | |
| 2528_D | ATAT<u>TCTAGA</u>ACCCCACCCAAATGAGGCAG | |
| 2528_E | AGCACGGTGACCCACAGCAG | |
| 2528_F | CCTGGCAGGAAGCCATGAAA | |
| 2528_G | AGAACGGGCGCGCGCCCATC | |
| 2528_H | CTTCTCCGTCAGCTCGCCGC | |
| 2604_A | ATCG<u>GGTACC</u>GCCAGGCTGTCTCCCACGAA | In-frame deletion of MXAN_2604 (*pixB*) |
| 2604_B | GTCATCCATCAGCACTGCGCGCGTCGT | |
| 2604_C | GCAGTGCTGATGGATGACAAGGACATC | |
| 2604_D | ATCG<u>TCTAGA</u>CGCAGGGCGGCGAGCAGCTT | |
| 2604_E | ATGACGTCTGCCTTCGTGCT | |
| 2604_F | CGGAACAGGTGGAACAGCTT | |
| 2604_G | AGCTCACGGTGACGGAGGAG | |
| 2604_H | TGGCGATGGCGCCGTAGATT | |
| 2649_A | ATCG<u>GGTACC</u>TGTCCGGCAAGGGCTGGCGT | In-frame deletion of MXAN_2649 |
| 2649_B | GGAAACGGATGCCTTCTGGCCACCCTT | |
| 2649_C | CAGAAGGCATCCGTTTCCGACCCTTGG | |
| 2649_D | ATCG<u>TCTAGA</u>CCATTTGCAGTCTTGTCATC | |
| 2649_E | ATCGTCATCTGGAACGACTA | |
| 2649_F | GATGTGGATGAAGAACTTCA | |
| 2649_G | TTCACGCCGTTGATTGAGAT | |
| 2649_H | TCCACCTCGTTGGGCAGGGA | |
| 2902_A | ATCG<u>GGTACC</u>TGGGAGAAGGGCTCGCCGCC | In-frame deletion of MXAN_2902 |
| 2902_B | CAAGCCGTGGGCGTAGGCGGACACAGG | |
| 2902_C | GCCTACGCCCACGGCTTGTGGAAAGGA | |
| 2902_D | ATCG<u>TCTAGA</u>CGGCCCTCGAACCGCAAGCT | |
| 2902_E | AGCAGGTCCACCGTCGCGAG | |
| 2902_F | CGCCGGGCATCATGTTGCAG | |
| 2902_G | TGCTGCTCCAAGGCGAGACG | |
| 2902_H | CCAGGAAGTAGAGCGCCAGC | |
| 3585_A | ATCG<u>GGTACC</u>TCGTAGTGGACCTTGACGGT | In-frame deletion of MXAN_3585 |
| 3585_B | ACGGTCGACCCGCTTGTTCTTCCGCTT | |
| 3585_C | AACAAGCGGGTCGACCGTCACGGCAAC | |
| 3585_D | ATCG<u>TCTAGA</u>GCCGCTGTCATCCAAGAGGC | |
| 3585_E | TGGTGCCGTCCACCAGCCGG | |
| 3585_F | AGCGCCGCCGCGCCGTAGAG | |
| 3585_G | CGACATCTACCTGAACAAGT | |

| | | |
|---|---|---|
| 3585_H | ATCATCTTGCGGTGACGATC | |
| 3721_A | ATCGGGTACCTTCGTCAACGACGCGGAGGC | In-frame deletion of MXAN_3721 |
| 3721_B | CTTCACCTGGCTCAACCCGGAACCTTC | |
| 3721_C | GGGTTGAGCCAGGTGAAGGTCGGCTGG | |
| 3721_D | ATCGTCTAGACCATTGCCAATGGCGAAGGC | |
| 3721_E | TCTTCCTGGGCAACATCCAC | |
| 3721_F | TCCCCGCGCCTGTCCGCCTT | |
| 3721_G | TTCCCACCCAGCTTCGGTGC | |
| 3721_H | GCGAAGCCGTCCCAGGGTAT | |
| SK68_A | GCGCGAATTCGGAAGTTCTTCCGGGTGT | In-frame deletion of MXAN_3778 |
| SK69_B | TCCGAACAAGACCGCCGCCTGATTCGA | |
| SK70_C | GCGGCGGTCTTGTTCGGAAGAGCCTGA | |
| SK71_D | GCGCTCTAGAAAGTTGGGTCCACCGCTG | |
| SK110_E | CACCTTCACCGCGTCCACGC | |
| SK111_F | GGGCTCCGCGCCGCAGAA | |
| SK112_G | TTCCCCCGGTTCCTCCCAAT | |
| SK113_H | ATGGAGGGCACCGTGTTGTG | |
| SK245_A | GCCGGGTACCCCGGACCTCATCAACGCCGGG | In-frame deletion of MXAN_3788 |
| SK246_B | CTATTCGGGCGGGAGCTCGCCGCGGGGTTCGTACA | |
| SK247_C | TTGTACGAACCCCGCGGCGAGCTCCCGCCCGAATAG | |
| SK248_D | GCCGTCTAGAGCACGGGCGTGAGCGCGC | |
| SK249_E | ACGCCTGCCTCCCTGGTACA | |
| SK250_F | GTGACGCCGCAGGGACTCAT | |
| SK251_G | CGCTCCACGAACTCCGCGATA | |
| SK252_H | CAGTCCCAACACCTCGACCCC | |
| 4328_A | ATCGGGTACCATCGCCCGGTGAGGTGACGC | In-frame deletion of MXAN_4328 |
| 4328_B | GCGGTCGACACGGCGCTCGTGAGGACG | |
| 4328_C | GAGCGCCGTGTCGACCGCGAGCGGGTG | |
| 4328_D | ATCGTCTAGAGCGCCCGCCTGCTTGCGGTG | |
| 4328_E | GAGGCGCTCCAGTACATCGC | |
| 4328_F | CCGCCAACATCCTGTCGCAG | |
| 4328_G | ACCTCCGTTTCGACAAGGTC | |
| 4328_H | GTCGTCCTGCGTCTCATAGG | |
| SK164_A | GCGCGGTACCGCTGGGTGACGTCATTCTCC | In-frame deletion of MXAN_4567 |
| SK165_B | CTATCGCTTGAAGCTGTCGCTGCTTCCCACACCCAC | |
| SK166_C | GTGGGTGTGGGAAGCAGCGACAGCTTCAAGCGATA | |
| SK167_D | GCGCTCTAGACTGATGCGCCGGCCTTCCGT | |
| SK197_E | CGGCTTCATTCCCATTCCCA | |
| SK198_F | GGACACCGAAGTGACGAACT | |
| SK199_G | GTGCGTATGACAACCAAGGC | |
| SK200_H | GTGGCTTCACCGAAATCGAC | |
| SK152_A | GCGCGAATTCCCCATCCGCCGGCCGGAGCA | In-frame deletion of MXAN_5615 |
| SK261_B1 | TCACGGCATCCGCGCGACCAGGGGGGAACTCAAGT | |
| SK262_C1 | AACTTGAGTTCCCCCCTGGTCGCGCGGATGCCGTGA | |
| SK155_D | GCGCTCTAGAGGGCGTGCTGGGGGGCATGA | |

| | | |
|---|---|---|
| SK172_E | CAGCTCGATGACCTTCACCAT | |
| SK173_F | GAGTACGCCATCGAGTCTTCC | |
| SK174_G | GCAAAGTGTCAGTCGCGTG | |
| SK175_H | GAGTGTTCATCGGCGGAGTAG | |
| SK237_A | GCCGGGTACCCCACGCCTCCTGGTTGCCCGC | In-frame deletion of MXAN_5655 |
| SK238_B | TCAGGGCAGGCGCGGGTGGTTTCTCCTCTTCTCCAC | |
| SK239_C | GTGGAGAAGAGGAGAAACCACCCGCGCCTGCCCTG | |
| SK240_D | GCCGTCTAGATTGCATCCTCCCTGGTGCGGC | |
| SK241_E | AGCACCGTGCCGAAGTCGTT | |
| SK242_F | TTGGGCAATGGCCCGTGTGA | |
| SK243_G | GACGCTCGGTGATGGTTCGC | |
| SK244_H | GGCGAATCTGGCAGGTGGTGA | |
| SK352_R | GCACCACTCCGCTTCATTCG | |
| 5707_A | ATCGGGTACCCAGGCACGAGGTGCCCCGAGA | In-frame deletion of MXAN_5707 |
| 5707_B | GCTCTGCACGCTCATGCTCGCGCCACC | |
| 5707_C | AGCATGAGCGTGCAGAGCGGCACGGCC | |
| 5707_D | ATCGTCTAGAAGTCAGGAGCAAATCCACGGC | |
| 5707_E | GAACTTCGGACGCGCACTCTA | |
| 5707_F | AGAGCGTCTGACAAGCGTGGA | |
| 5707_G | GACTCGATGAGCGACAAGGCC | |
| 5707_H | GGCCTGGAGCACCTCGCCCTT | |
| SK036_A | GCGCGGTACCAGGCTCGCGATGTGAATG | In-frame deletion of MXAN_5804 |
| SK037_B | GCCGAAGAGGGACGCTGTCCCGCTCAT | |
| SK038_C | ACAGCGTCCCTCTTCGGCCGCCGCTGA | |
| SK039_D | GCGCTCTAGACTGGCGGCGGAGATGCTG | |
| SK050_ E | GGGACGACGGGCTACACCTGTTGC | |
| SK051_F | TTCTCGTATCGCGCCTATGTGGAGT | |
| SK052_G | AGTCCGACGCCGAAGCAGGC | |
| SK053_H | CACGCTCACGCCCAGGCTCTCTT | |
| SK72_A | GCGCGAATTCGTTCCTGGACCTGTTCAT | In-frame deletion of MXAN_6013 |
| SK73_B | CTTCTTCAGGGCCGGCAGGCTCTGCAT | |
| SK74_C | CTGCCGGCCCTGAAGAAGAAGCCCTGA | |
| SK75_D | GCGCTCTAGAGGGCAGCGGCCCACCGCG | |
| SK089_E | CGAGGCTGCGGACGGTGA | |
| SK090_F | TCAGGGCCAGCGTGCGCT | |
| SK091_G | AGCCTGGACGAGCTGCGTCT | |
| SK092_H | AATTCTCCCAGATGACGGAGCTGCG | |
| 6605_A | ATCGGAATTCACTCGCCGCTCTTGCGCTGGT | In-frame deletion of MXAN_6605 |
| 6605_B | CTTCTTCACGTCCGTGGTGCGCCTGCC | |
| 6605_C | ACCACGGACGTGAAGAAGGCGCTGGGC | |
| 6605_D | ATCGTCTAGACAGTAGAAGCCGTCCAACCTG | |
| 6605_E | TGTGAACCACGCCAGGCTGCC | |
| 6605_F | GCAGTGCCCGTCCTCGCGCTT | |
| 6605_G | AAGGGCTTCCCCAAGTGCGCC | |
| 6605_H | TTGAGCTCGCGGTTGAGACCG | |

| | | |
|---|---|---|
| SK148_A | GCGCGAATTCACCACAACCACTGCCCCACC | In-frame deletion of MXAN_6957 |
| SK149_B | TCACGGGCGCACCTCCGCTCCTGGAGCCGTCATCAT | |
| SK150_C | ATGATGACGGCTCCAGGAGCGGAGGTGCGCCCGTG | |
| SK151_D | GCGCTCTAGAACCGATGAGGATGCGCAGCG | |
| SK168_E | ATTCCGACGCACAACCTCAA | |
| SK169_F | CCGGTGGACCTGAAGGATTT | |
| SK170_G | CCACGAGCGAATCCTGGTT | |
| SK171_H | CGCTGTAGAGGATGCGATGA | |
| SK76_A | GCGCGAATTCGGTCGTCTTCGAGCTGTC | In-frame deletion of MXAN_7024 |
| SK77_B | TGAGGCGCGGTTGGGAGGAGGGCTCAT | |
| SK78_C | CCTCCCAACCGCGCCTCAGGTTTCTGA | |
| SK79_D | GCGCTCTAGATTGTTCGTGCCCGTGTAG | |
| SK106_E | GCGAGGTGGTGCTGAAGC | |
| SK107_F | CACGAACGGTGGATGTAGAGC | |
| SK108_G | CCGACGACTTCCTGCCCAAG | |
| SK109_H | GGAACCAGTCGCCCCAGAAC | |
| SK128 | GCGCTCTAGAAGTACCGCACGAGGCCAG | Complementation of Δ*pixA* |
| SK129 | GCGCAAGCTTCTACTTGATGCGCAAGAGCTG | |
| SK384 | GCCGAAGCTTTCACTTGTCGTCGTCGTCCTTGTAGTC CTTGATGCGCAAGAGCTG | *pixA*-FLAG |
| SK385 | CTTGTTATGAACCCGGGTCCAGAGGACAAG**GCC**CAG CACCCCCGCGTCCCGGCCGTGCTGAG | R9A point mutation in *pixA* |
| SK386 | CTCAGCACGGCCGGGACGCGGGGGTGCTG**GGC**CTT GTCCTCTGGACCCGGGTTCATAACAAG | |
| SK189 | GGTGATGACG**GAG**CTCTACATGC | D180E point mutation in *pixA* |
| SK190 | GCATGTAGAG**CTC**CGTCATCACC | |
| SK191 | GGTGATGACG**AAC**CTCTACATGC | D180N point mutation in *pixA* |
| SK192 | GCATGTAGAG**GTT**CGTCATCACC | |
| Pnat 2604 XbaI fw | ATCGTCTAGAGGCACCTCCCGACTCCCACC | Complementation of Δ*pixB* |
| 2604 KpnI rev | ATCGGGTACCTCATGCCGCGATGTCCTTGT | |
| 2604 XbaI fw | ATCGTCTAGAATGACCGTGACGACGCGCGC | |
| 2604 FLAG Rw | GCGAAGCTTTCACTTGTCGTCGTCGTCCTTGTAGTCT GCCGCGATGTCCTT | *pixB*-FLAG |
| R121A PilZ1 2604 fw | GAATTGCTGGAACTG**GCG**GAGTCCGAGCGCCGC | R121A point mutation in *pixB* |
| R121A PilZ1 2604 rev | GCGGCGCTCGGACTC**CGC**CAGTTCCAGCAATTC | |
| R331A 2604 fw | ACCACCGC**GCG**AACAGCCTG | R331A point mutation in *pixB* |
| R331A 2604 rev | CAGGCTGTT**CGC**GCGGTGGT | |

| | | |
|---|---|---|
| 2604 GNAT-FLAG (451) rev | gcgaagctttcacttgtcgtcgtcgtccttgtagtcACACCGTGCCTTGAT | Deletion of AT domain in *pixB* |
| pSK143 | GCCGTCTAGACAGCGGCTGGGCAATGGCAT | Deletion of both PilZ domains in *pixB* |
| SK122 | gcgcCATATGaacccgggtccagaggac | Purification of *pixA*-His$_6$ |
| SK123 | gcgcCTCGAGcttgatgcgcaagagctgc | |
| 2604 NdeI fw | ATCGCATATGATGACCGTGACGACGCGCGC | Purification of *pixB*-His$_6$ |
| 2604 HindIII -stop rev | ATCGAAGCTTTGCCGCGATGTCCTTGTCAT | |

* Restriction sites are underlined and mutations introduced by site-directed mutagenesis are in bold.
